# The *Fox* gene repertoire in the annelid *Owenia fusiformis* reveals multiple expansions of the *foxQ2* class in Spiralia

**DOI:** 10.1101/2022.03.02.482670

**Authors:** Océane Seudre, Francisco Manuel Martín-Zamora, Valentina Rapisarda, Allan M. Carrillo-Baltodano, José M. Martín-Durán

## Abstract

*Fox* genes are a large and conserved family of transcription factors involved in many key biological processes, including embryogenesis and body patterning. Although the role of *Fox* genes has been studied in an array of model systems, comprehensive comparative studies in Spiralia—a large clade of invertebrate animals including molluscs and annelids—are scarce but much needed to better understand the evolutionary history of this gene family. Here, we reconstruct and functionally characterise the *Fox* gene complement in the annelid *Owenia fusiformis*, a slow evolving species and member of the sister group to all remaining annelids. The genome of *O. fusiformis* contains at least a single ortholog of each of the 23 *Fox* gene classes that are ancestral to Bilateria, except for *foxE* and *foxI*. Temporal and spatial expression dynamics reveal a conserved role of *Fox* genes in gut formation (*foxA*), mesoderm patterning (*foxF, foxL1*, *foxC*, *foxH*) and cilia formation (*foxJ1*) in Annelida and Spiralia. Moreover, we uncover an ancestral expansion of *foxQ2* genes in Spiralia, represented by 11 paralogs in *O. fusiformis*. Notably, although all *foxQ2* copies have apical expression in *O. fusiformis,* they show variable spatial domains and staggered temporal activation, which suggest cooperation and sub-functionalisation among *foxQ2* genes for the development of apical fates in this annelid. Altogether, our study informs the evolution and developmental roles of *Fox* genes in Annelida and Spiralia generally, providing the basis to explore how regulatory changes in *Fox* gene expression might have contributed to developmental and morphological diversification in Spiralia.

**Significance statement:** The role of *Fox* genes, a group of DNA-binding proteins required for the formation of many animal organs, is poorly understood in invertebrate groups such as molluscs and annelids. Here, by studying the genome and embryogenesis of the annelid *Owenia fusiformis*, we demonstrate that *Fox* genes are involved in the development of the gut, muscles, cilia, and nervous system. Importantly, we find that a group of *Fox* genes (referred to as *foxQ2*) expressed in the anterior end of most animals has more copies in annelids and molluscs than in other invertebrate groups like insects and sea stars. Together, our findings clarify the evolution of *Fox* genes and their contribution to the diversity of forms and organs found in marine invertebrates.

## Introduction

Forkhead box-containing proteins (i.e., *Fox* genes) form one of the largest families of transcription factors in animals, displaying a remarkable functional diversity in many morphogenetic processes (Carlsson & Mahlapuu 2002; Hannenhalli & Kaestner 2009; Kaufmann & Knöchel 1996). *Fox* genes are characterised by a conserved DNA-binding domain of approximately 100 amino acids—the Forkhead or winged helix domain—that folds into two stereotypical large loops or “wings” (Clark et al. 1993; Li & Tucker 1993). Since the discovery of the Forkhead domain in the fruit fly *Drosophila melanogaster* (Weigel et al. 1989), *Fox* genes have been studied in a wide range of traditional developmental systems, mostly vertebrates (Mazet et al. 2003; Hannenhalli & Kaestner 2009; Lee & Frasch 2004; Jackson et al. 2010; Golson & Kaestner 2016). The initial description of 15 *Fox* gene classes in chordates, each identified by a letter (Kaestner et al. 2000), and the establishment of a unified nomenclature facilitated phylogenetic analyses and comparisons with other major invertebrate clades, such as hemichordates (Fritzenwanker et al. 2014), molluscs (Mei *et al*., 2014; Wu *et al*., 2020), platyhelminthes (Pascual-Carreras et al. 2021), panarthropods (Schomburg et al. 2022), cnidarians (Magie et al. 2005), and sponges (Larroux et al. 2006), as well as animal outgroups (Larroux et al. 2008), ultimately uncovering a complex evolutionary history for this large family of transcription factors. Today, *Fox* genes are classified into 26 classes belonging to two major clades (Larroux et al. 2008; Mazet et al. 2003; Hannenhalli & Kaestner 2009; Benayoun et al. 2011; Kaestner et al. 2000; Pascual-Carreras et al. 2021), where ancestral duplication events (e.g., the former class *foxQ* split into *foxQ1* and *foxQ2*, *foxN* into *foxN1/4* and *foxN2/3*, *foxL* into *foxL1* and *foxL2*, and *foxJ* into *foxJ1* and *foxJ2/3*), gene innovations (e.g., *foxR* and *foxS* are unique of vertebrates, *foxT* is a novelty of panarthropods), expansions and losses (e.g., *foxAB* in vertebrates, *foxQ2* in tetrapods, and *foxAB*, *foxE*, *foxH*, *foxI* in arthropods) are common (Paps et al. 2012; Schomburg et al. 2022; Mazet et al. 2003; Wotton & Shimeld 2006; Tu et al. 2006). Moreover, genomic comparisons also uncovered conserved syntenic linkage for some of the classes, such as *foxL1*-*foxC*-*foxF*-*foxQ1*, in phylogenetically distant lineages of insects, chordates and spiralians (Mazet et al. 2006; Wotton & Shimeld 2006, 2011; Wotton et al. 2008; Shimeld, Boyle, et al. 2010). Yet a comprehensive characterisation of *Fox* genes is still lacking in most major animal groups, most notably in members of Spiralia, one of the two main clades of protostomian animals that comprises nearly half of the extant major metazoan groups, including molluscs and annelids (Marlétaz et al. 2019). Therefore, the evolutionary history and developmental roles of this conserved family of transcription factors are still unclear at key nodes of the animal tree of life.

*Fox* genes typically show tissue-specific expression patterns and play an important role in cell-type determination and differentiation (Hannenhalli & Kaestner 2009; Jackson et al. 2010). Functional studies in human, mouse, zebrafish and *D. melanogaster* have revealed an array of functions of *Fox* genes in early development, such as axial patterning, germ layer specification, and organogenesis (reviewed in Carlsson & Mahlapuu, 2002). In Spiralia, however, studies on the function of *Fox* genes are scarce and mostly focused on certain classes, with just a handful of studies encompassing more than one major spiralian clade (Supplementary Table 1 and references therein). For example, *foxA* is consistently expressed in the developing foregut in many spiralians, including annelids, brachiopods, phoronids, and bryozoans (Arenas-Mena 2006; Martín-Durán et al. 2016; Andrikou et al. 2019; Vellutini et al. 2017; Adler et al. 2014; Boyle & Seaver 2008, 2010; Kwak et al. 2018; Kostyuchenko et al. 2019) and *foxJ1*, *foxQ2* and *foxG* are expressed in larval specific tissues in the annelid *Platynereis dumerilii*, the brachiopod *Terebratalia transversa* and the phoronid *Phoronopsis harmerii* (Marlow et al. 2014; Santagata et al. 2012; Gąsiorowski & Hejnol 2020). Similarly, the clustered classes *foxC*, *foxL1* and *foxF* show mesodermal expression in all spiralian species studied to date, suggesting that coordinated activation of these *Fox* genes in a common germ layer might have contributed to the maintenance of their genetic linkage (Shimeld, Boyle, et al. 2010; Passamaneck et al. 2015; Martín-Durán et al. 2016). Other *Fox* gene classes, however, have only been studied in individual species, which prevents inferring an ancestral role for these genes in Spiralia. For example, *foxL2* is a regulator of ovarian differentiation and development in molluscs (Liu et al. 2012; Li et al. 2016; Mei, Fei, Jun, Li, Yang, Xu, Liu, Que, Li & Zhang 2014; Teaniniuraitemoana et al. 2015), *foxB* is expressed during late mesoderm development in the leech *Helobdella austinensis* (Kwak et al. 2018), *foxO* controls tissue regeneration and cell death in the planarian *Schmidtea mediterranea* (Pascual-Carreras et al. 2021) and *foxK1* is involved in ectodermal regeneration in that same planarian species (Coronel-córdoba et al. 2022). Consequently, the repertoire and developmental functions of most *Fox* genes remain largely unexplored in Spiralia, and thus its study is not only important to discern the evolution of this gene family in animals, but also the contribution of these developmental regulators to the diversification of body plans and embryonic modes in this major animal group.

Here, we mine the genome of the annelid *Owenia fusiformis* (Liang et al. 2022), a member of Oweniidae and sister group to all remaining annelids (Rouse et al. 2022), to infer the ancestral *Fox* gene complement to Annelida, one of the most species-rich and morphologically diverse groups within Spiralia. Temporal and spatial gene expression analyses offer insights into the potential role of some of the *Fox* gene classes in this annelid, uncovering conserved and putative new roles for some of the *Fox* classes. Moreover, our study reveals that the *foxQ2* class is largely expanded in Spiralia, with the paralogs being consistently expressed in apical territories and exhibiting signs of possible sub-functionalisation in *O. fusiformis*. Altogether, our study informs the evolution of the *Fox* gene family in Annelida, providing valuable data to reconstruct the evolution and developmental roles of these genes in Spiralia and Metazoa.

## Results

### The *Fox* gene complement in *O. fusiformis*

To characterise the *Fox* gene complement in the annelid *O. fusiformis,* we searched for annotated gene models containing the conserved Forkhead DNA-binding domain in its reference genome (Liang et al. 2022). We obtained a total of 34 putative *Fox* genes, which is a number considerably higher than those previously reported in other Spiralian species, in which the number of *Fox* genes ranges from 21 to 26 genes (Wu et al. 2020; Mei, Fei, Jun, Li, Yang, Xu, Liu, Que, Li & Zhang 2014; Pascual-Carreras et al. 2021). To assign the orthology of each of the *O. fusiformis Fox* genes, we applied maximum likelihood and Bayesian phylogenetic tree inference, obtaining strongly supported orthologs for 20 of the 25 classes of *Fox* genes, both in clade I and clade II (Figure 1A, B; Supplementary Figures 1–4). Only the two ancestral bilaterian *Fox* gene classes *foxE* an *foxI* are missing in *O. fusiformis*, which have been however reported in other spiralian lineages (Figure 1C) (Wu et al. 2020; Mei, Fei, Jun, Li, Yang, Xu, Liu, Que, Li & Zhang 2014). In addition, *O. fusiformis* has a fast evolving ortholog that groups with low node support with other fast evolving Forkhead-containing genes collectively termed *foxY*, which were first described in *S. purpuratus* (Tu et al. 2006) and later identified in the spiralians *Crassostrea gigas*, *Lottia gigantea*, *Mizuhopecten yessoensis* and *Capitella teleta* (Mei, Fei, Jun, Li, Yang, Xu, Liu, Que, Li & Zhang 2014; Wu et al. 2020) (Figure 1C; Supplementary Figure 1, 3). While most *Fox* gene classes have a single ortholog in *O. fusiformis*, we also detected duplication events in *foxAB*, *foxN1/4* and *foxN2/3* classes, as well as a large expansion in *foxQ2*, which has 11 paralogs (Figure 1A–C; Supplementary Figures 1–4). Together, our findings demonstrate that *O. fusiformis* has a more conservative Fox gene complement than all other spiralian lineages previously characterised (Figure 1C), thus helping to infer an ancestral *Fox* repertoire and lineage-specific losses in Spiralia.

**Fig. 1.**
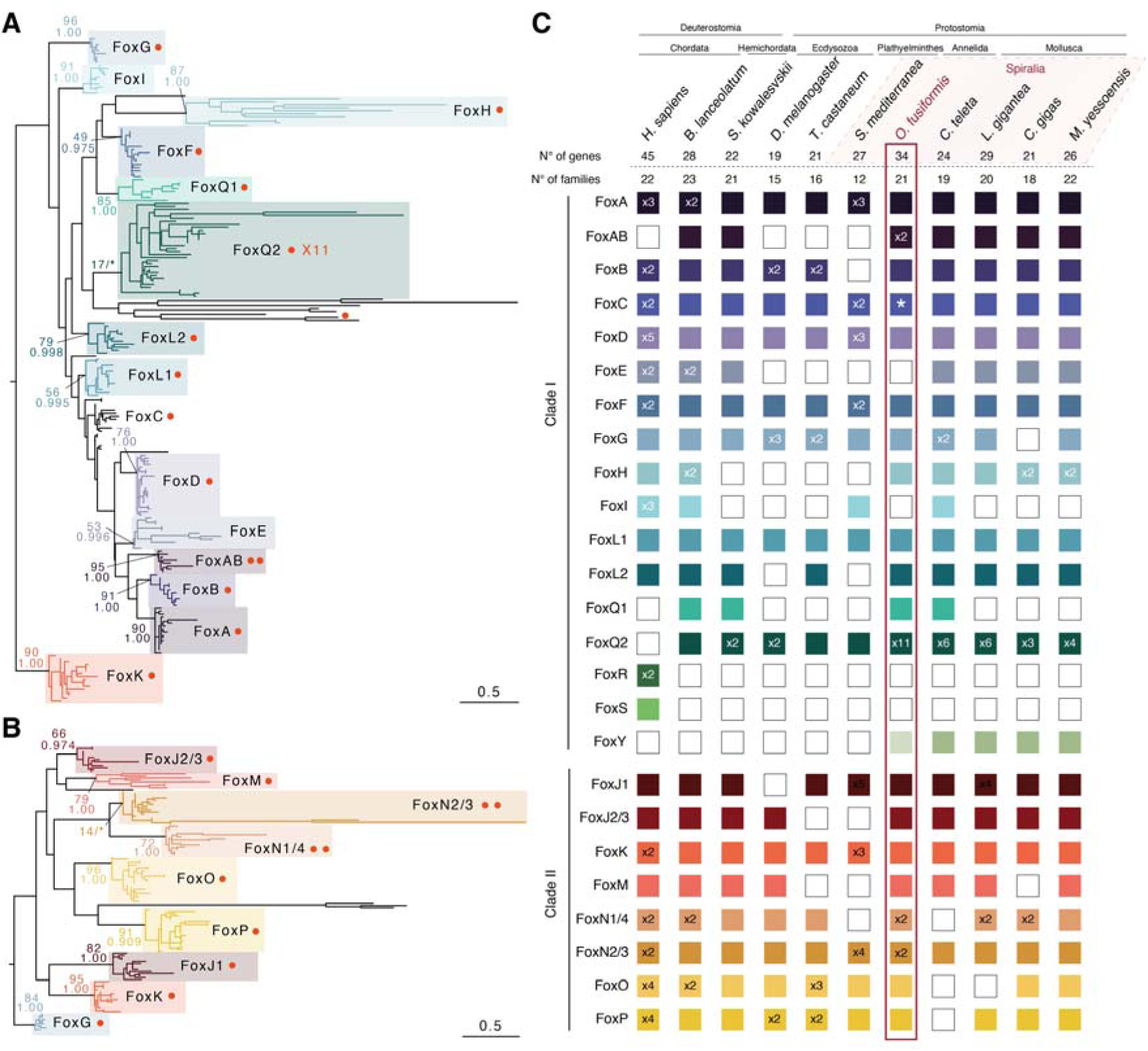
The *Fox* gene complement in *O. fusiformis* and selected metazoan clades. (**A**, **B**) Orthology assignment of *Fox* genes in Clade I (**A**) and Clade II (**B**) in *O. fusiformis* and 18 other metazoan taxa, using the *foxK* and *foxG* classes as outgroups for Clade I and Clade II, respectively. Tree topologies are based on maximum likelihood reconstruction and node supports indicate both bootstrap values and posterior probabilities at key nodes. Coloured boxes indicated each *Fox* gene class and red dots indicate the presence and number of copies in *O. fusiformis* genome. Scale bars indicate the number of amino acids substitutions per site alongside the branches. (**C**) Summary of *Fox* gene repertoires in *O. fusiformis* and other metazoan species. Coloured boxes indicate the presence of an ortholog and numbers inside specify the number of paralogs per class and species. The asterisk in the *foxC* class indicates that the presence of a *foxC* gene in *O. fusiformis* is based on a previous report.

### Genomic architecture and chromosomal linkage of *Fox* genes in *O. fusiformis*

*Fox* genes belonging to clade I and clade II additionally differ in whether they lack or contain introns at conserved sites of the Forkhead domain, respectively (Larroux et al. 2008). To further characterise the *Fox* gene complement and assess this rule in *O. fusiformis*, we reconstructed the domain architecture and exon-intron composition of all 34 *Fox* genes in this annelid. *Fox* genes in *O. fusiformis* show diverse gene architectures, with lengths ranging from 339 bp (*foxN1/4-b*) to 2,223 bp (*foxL1*), and all but *foxQ2-10* contain an intact Forkhead domain (Figure 2A). One *Fox* gene—*foxQ2-10*— shows additional protein domains, namely a DNA translocase domain (Figure 2A), which might indicate that it has acquired new protein functions. Exon numbers in the *Fox* genes of *O. fusiformis* range from one to 12, and most genes follow the clade I and clade II distinction based on the number of introns, apart from *foxF*, *foxAB-b, foxL2*, *foxY* and *foxQ2-6*, which contain introns albeit they belong to clade I and might thus represent independent intron gains (Figure 2A; Supplementary Table 2). Altogether, these findings reinforce the previous observation that *O. fusiformis* contains a well conserved *Fox* gene repertoire, while they highlight as well that a handful of *Fox* gene classes and paralogs might have experience faster rates of molecular evolution.

**Fig. 2.**
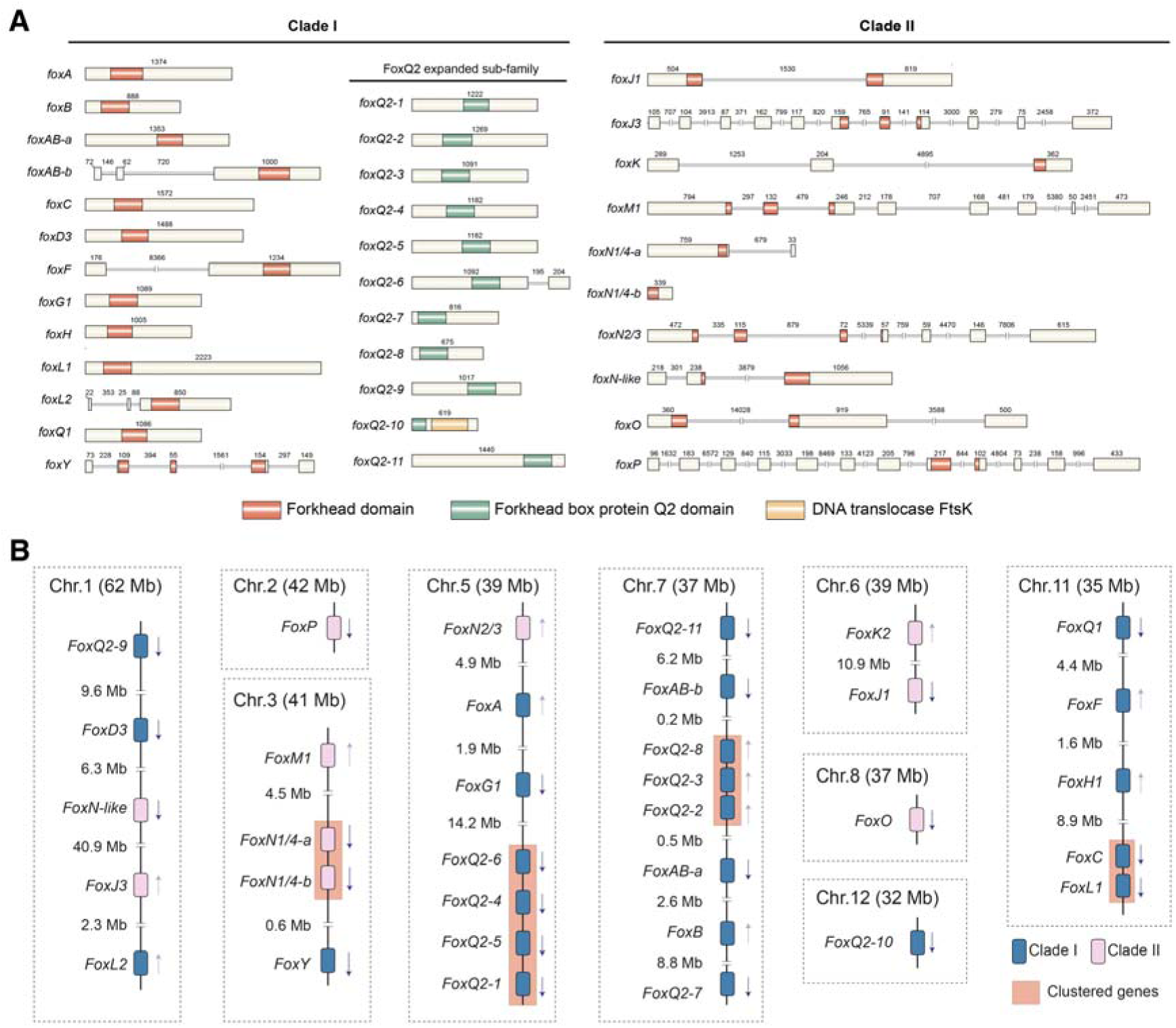
The genomic architecture and linkage of *Fox* genes in *O. fusiformis*. (**A**) Schematic representation of the protein domain composition and exon-intron structure for each *Fox* gene in *O. fusiformis*. Beige boxes indicate coding regions and double black lines represent introns (double oblique lines mean the scale is not proportional and number above exons and introns indicate their length in base pairs). Red boxes depict the location of the Forkhead domain (PF00250), green boxes show the FoxQ2 specific domain (CD20035), and an orange box highlights the DNA translocase FtsK domain in *foxQ2-10*. (**B**) Schematic drawings of the genomic organisation of the *Fox* genes in *O. fusiformis*. For each chromosome (Chr; chromosome size in brackets and in Mb), numbers between genes indicate major intergenic distances in Mb and arrows indicate the transcriptional orientation of each gene. *Fox* genes belonging to Clade I are in blue and *Fox* genes belonging to Clade II are in pink. Clustered genes are highlighted with an orange box.

Certain *Fox* gene classes (e.g., *foxL1*, *foxC*, *foxF*, and *foxQ1*) show conserved chromosomal linkage across phylogenetically distant bilaterian taxa (Shimeld, Boyle, et al. 2010; Shimeld, Degnan, et al. 2010; Mazet et al. 2006). To assess whether this feature was also retained in *O. fusiformis*, we study the chromosomal location and microsyntenic relationship of the 34 *Fox* genes in this annelid. *Fox* genes are spread across nine of the 12 chromosomes of *O. fusiformis*, namely chromosomes 1, 2, 3, 5, 6, 7, 8, 11, and 12 (Figure 2B). While *foxF*, *foxC*, *foxL1*, and *foxQ1* are located on the same chromosome—chromosome 11—only *foxC* and *foxL1* show evidence of a tight linkage, with a high number of genes (>1,000) lying between f*oxQ1* and *foxF* as well as between f*oxF* and *foxC* (Figure 2B). In addition, we observed evidence of tandem duplications in some of the *Fox* classes that exhibit expansions, in particular *foxN1/4* (on chromosome 3) and two clusters of multiple copies of *foxQ2* genes in chromosome 5 and 7 (Figure 2B). Therefore, even though the ancestral bilaterian chromosomal linkage is overall conserved in *O. fusiformis* (Liang et al. 2022), the ancestral microsyntenic relationships observed among certain *Fox* genes is lost in this annelid species.

### The expression dynamics of the *Fox* gene complement in *O. fusiformis*

To investigate the expression dynamics of the *Fox* genes in *O. fusiformis* and relate each of these genes to major morphogenetic events during the life cycle of this annelid, we used available stage-specific RNA-seq data covering 14 developmental time points, from the unfertilized oocyte to the juvenile stage (Liang et al. 2022). In *O. fusiformis*, the temporal expression dynamics of the *Fox* genes seem to correlate with their assignment to clade I and clade II, because most clade II *Fox* genes are expressed maternally and during the early cleavage stages (i.e., up to the 8-cell stage), while clade I *Fox* genes tend to show short peaks of expression at single developmental stages, from the 32-cell stage onwards (Figure 3A). Clade II genes that escape this trend are *foxP*, expressed at the juvenile stage, *foxN2/3*, which peaks during gastrulation, and *foxJ1*, *foxK2*, and *foxO* that are expressed during larval development (Figure 3A). We confirmed these temporal expression profiles for the gene *foxJ1*, which is involved in cilia development during the embryogenesis of *O. fusiformis*, becoming expressed in the cells forming the apical organ and ciliated band in the larva of this annelid species (Figure 3B). Notably, a set of *Fox* genes belonging to clade I (*foxAB-a*, *foxG1*, and many *foxQ2* paralogs, see below) are finely expressed at the time of the specification of the embryonic organiser and the establishment of the axial identities in *O. fusiformis*, as well as during gastrulation (i.e.*, foxAB-b, foxH1, foxA*) (Seudre et al. 2022; Carrillo-Baltodano et al. 2021) (Figure 3A). Other genes from the same clade become expressed later during embryogenesis and are probably associated with either organogenesis and larval development (i.e., *foxD3*, *foxF*, *foxL1*, *foxC*) or juvenile metamorphosis (i.e., *foxB, foxL2, foxQ1*) (Figure 3A, C). Together, these data support an embryonic role for most *Fox* genes in *O. fusiformis*, revealing diverse expression dynamics that correlate with crucial cell-type specification and morphogenetic events.

**Fig. 3.**
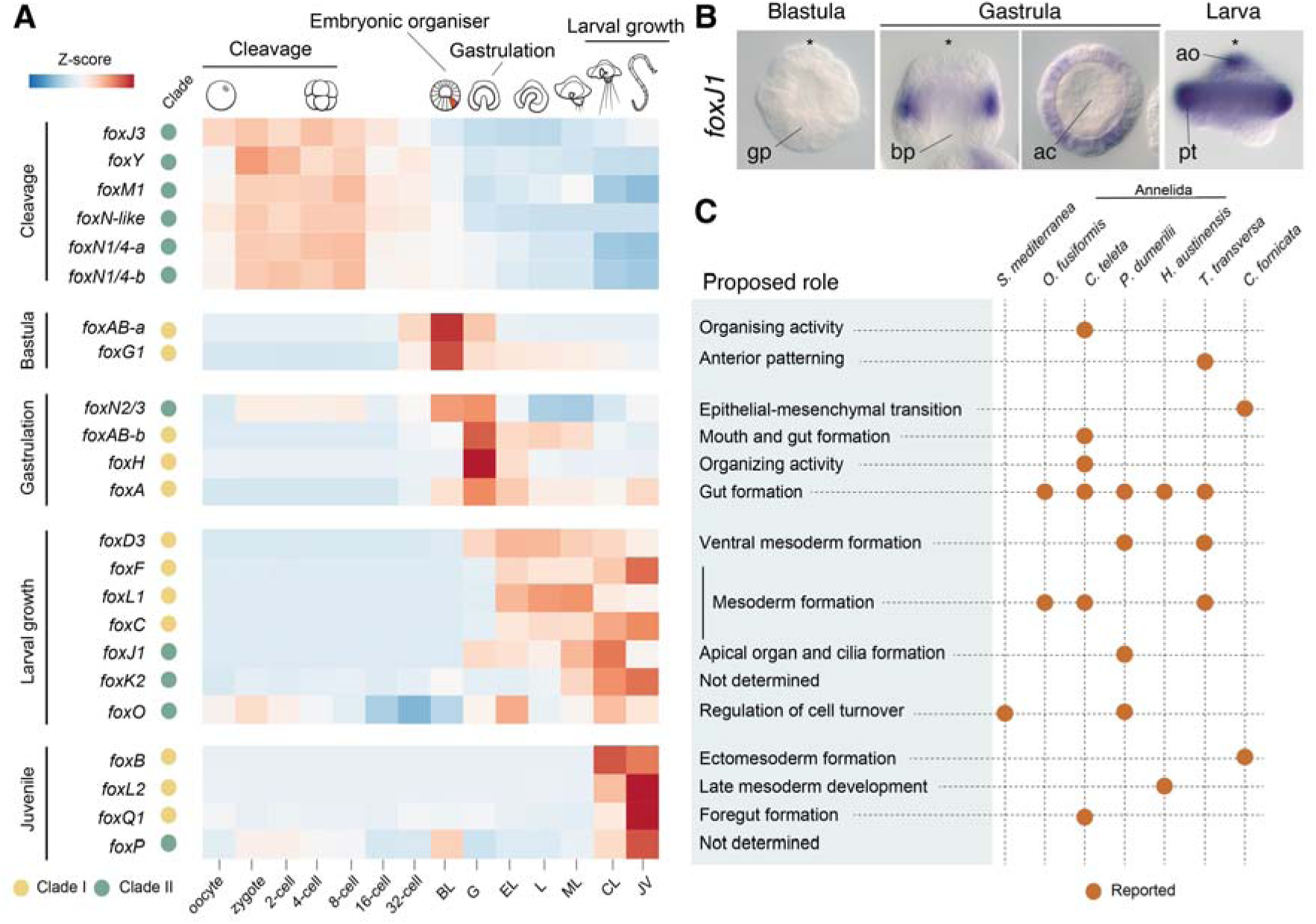
Temporal dynamics of expression of the *Fox* genes in *O. fusiformis* and their role in Spiralia. (**A**) Heatmap of expression of all *Fox* genes (except for the *foxQ2* family, which is shown in Figure 5) throughout development in *O. fusiformis*. Colours show the normalized z-score value of expression, with red blocks indicating high levels of expression and blue blocks indicating low levels of expression. The x-axis shows the developmental time points (BL: Blastula; G: Gastrula; EL: Elongation; L: Larva; ML: Mature larva; CL: Competent Larva; and JV: Juvenile). The second column from the left with yellow and green dots indicates the membership of each *Fox* gene to either Clade I or Clade II. Genes are ordered vertically following their timing of expression: maternal and during early cleavage; at the time of specification of the organiser cell (5 hours post fertilisation); during gastrulation; during larval growth; and at the juvenile stage. (**B)** Whole mount *in situ* hybridisation of *foxJ1* gene during the development of *O. fusiformis*, at the blastula (lateral view), gastrula (lateral to the left and ventral to the right) and larval stage (lateral view). Consistent with the temporal expression data, *foxJ1* starts to be expressed at the putative prototroch precursor cells at the gastrula stage and it is later detected in the ciliated cells of the larva (apical organ and prototroch). Asterisks mark the apical/anterior pole. ac, archenteron; ao, apical organ; bp, blastopore; gp, gastral plate; pt, prototroch. (**C**) Table summarising the current knowledge of the developmental roles of *Fox* genes in Spiralia (from Supplementary Table 1). For each gene, described developmental roles are in the light green box to the left and the associated species are shown with a red dot on the corresponding horizontal line on the right.

### Owenia fusiformis *has an expanded* foxQ2 *class*

Previous studies indicated that an ancestral duplication in the *foxQ2* class occurred at least in the last common bilaterian ancestor, and maybe even predated the cnidarian-bilaterian split (Fritzenwanker et al. 2014; Pascual-Carreras et al. 2021; Chevalier et al. 2006), which was followed by further duplications of this *Fox* gene class in some spiralian and deuterostomian lineages (Wu et al. 2020; Mei, Fei, Jun, Li, Yang, Xu, Liu, Que, Li & Zhang 2014). This observation agrees with the large expansion of 11 *foxQ2* paralogs that we identified in the annelid *O. fusiformis* (Figure 1A, C). Moreover, the presence of an EH-i-like Groucho binding motif in some *foxQ2* orthologs and its variable C- and N-terminal position with respect to the Forkhead and FoxQ2 domain has been used to subdivide the *foxQ2* class in *foxQ2-C* and *foxQ2-N*, or even to define a new sub-class named *foxQD* (Fritzenwanker et al. 2014; Pascual-Carreras et al. 2021). To clarify the evolution of this *Fox* gene class, and how the expansions of *foxQ2* genes occurred in *O. fusiformis* and spiralians generally, we mined available databases and the genomes of seven annelid species in search for genes with complete FoxQ2 domains, which we then used for phylogenetic reconstruction and the identification of EH-i-like motifs. While the general orthology of all identified *foxQ2* genes was robustly supported (Figure 4A), we did not recover two separate monophyletic clades with EH-i-like motifs in either the C- or the N-terminal end. Instead, *foxQ2* orthologs with an EH-i-like motif at the C-terminal end (Figure 4B) appear to form a relatively well supported monophyletic clade, comprising sequences of both cnidarians and most bilaterian groups, often as single copy genes (yet the annelids *O. fusiformis*, *Dimorphilus gyrociliatus* and *Helobdella robusta* have two paralogs). The rest of the *foxQ2* sequences lack an EH-i-like motif and probably represent more or less divergent copies. Among these fast-evolving *foxQ2* copies we found lineage-restricted expansions, such as those of *O. fusiformis*—for which the phylogenetic relationship between paralogs correlate well with their genomic linkage—and the vestimentiferan annelids *Lamellibrachia luymesi* and *Paraescarpia echinospica*, which have a group of *foxQ2* genes that independently aquired an EH-i-like motif on the N-terminal end (Figure 4A). Together, our findings corroborate previous analyses revealing a complex evolutionary history for the *foxQ2* gene class, probably dominated by fast rates of molecular evolution and/or frequent independent events of gene duplication in both cnidarian and bilaterian lineages, specially among spiralians, as well as its complete loss in tetrapods.

**Fig. 4.**
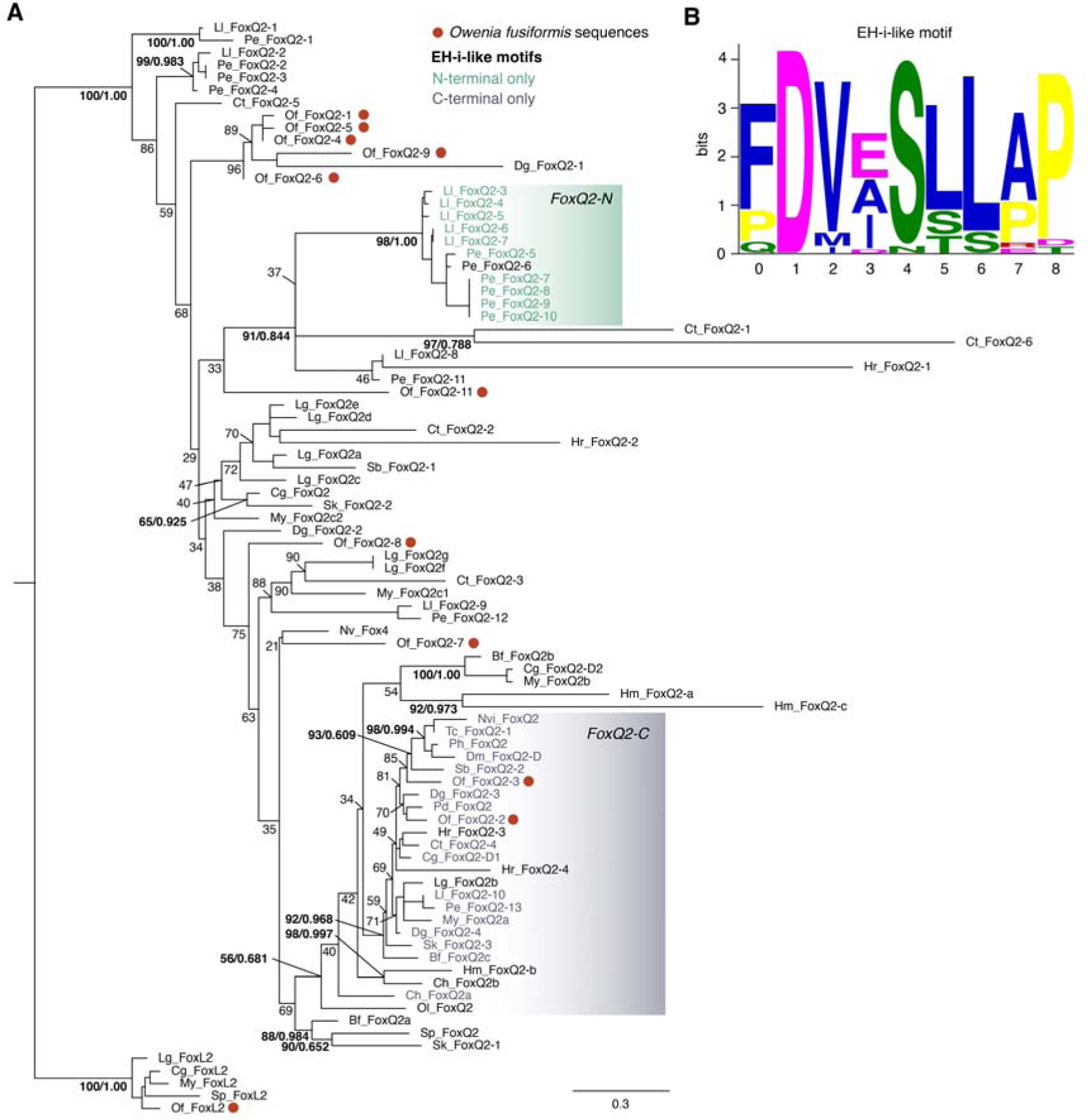
Phylogenetic analysis of the *foxQ2* family. (**A**) Maximum likelihood tree topology of the *foxQ2* class with *foxL2* class as outgroup. Sequence names are coloured based on the position of the EH-i-like motif (shown in (**B**)) with respect to the FoxQ2 box (sequences with a C-terminal position of the EH-I-like motif are in gray; sequences with an N-terminal position of the EH-I-like motif are in green; sequences lacking the EH-i-like motif are in black). The gray box highlights the monophyletic clade largely containing sequences of both cnidarian and bilaterian lineages with a C-terminal EH-i-like motif. Red dots indicate *O. fusiformis foxQ2* paralogs. Only sequences with a fully intact FoxQ2 domain (accession number CD20035) are included in the analysis (e.g., *foxQ2-10* from *O. fusiformis* is not included). (**B**) Sequence logo of the EH-i-like motif we identified.

### The developmental expression of *foxQ2* paralogs in *O. fusiformis*

To assess the potential functional implications of the expansions of *foxQ2* genes in spiralians (Figure 5A), we first compared the developmental expression profiles of *foxQ2* genes in *O. fusiformis* and three other spiralian species, namely the molluscs *Crassostrea gigas* and *Mizuhopecten yessoenssis* and the annelid *Capitella teleta* (Figure 5B–E). In the pacific oyster *C. gigas*, the three *foxQ2* paralogs show distinct temporal patterns of expression, with *foxQ2a* peaking at the trochophore stage, *foxQ2b* showing highest expression during gastrulation, and *foxQ2c* being expressed maternally and during the early cleavage stages (Figure 5B). Similarly, *foxQ2* genes in the scallop *M. yessoensis* display distinct temporal peaks of expression, with *foxQ2a* having a maximum at the blastula stage, *foxQ2c1* being strongly expressed during gastrulation, and both *foxQ2b* and *foxQ2c2* peaking at larval stages (Figure 5C). The six *foxQ2* paralogs of *C. teleta* display four different temporal patterns of expression: *foxQ2-1* is expressed in the oocyte*, foxQ2-2* and *foxQ2-3* peak during early cleavage, *foxQ2-4* is expressed at the blastula stage, and *foxQ2-5* and *foxQ2-6* show higher expression values during gastrulation (Figure 5D). Consistently, *foxQ2* paralogs are also expressed at distinct developmental stages in *O. fusiformis*, with *foxQ2-11* expressed first maternally and during early cleavage, *foxQ2-1, foxQ2-4, foxQ2-5, foxQ2-6, foxQ2-9, foxQ2-10* showing a peak of expression at the early blastula stage, *foxQ2-2* and *foxQ2-3* exclusively expressed at the time of the specification of the embryonic organiser at 5 hours post fertilisation and their expression being under control of the ERK1/2 signalling (Seudre et al. 2022), and finally both *foxQ2-7* and *foxQ2-8* coming up at larval stages (Figure 5E). Notably, spiralian *foxQ2* orthologs peaking at the blastula stage (i.e., *foxQ2-2* and *foxQ2-3* in *O. fusiformis*, *foxQ2-4* in *C. teleta*, *foxQ2a* in *M. yessoensis*, and *foxQ2-b* in *C. gigas*) belong to the strongly supported clade of *foxQ2* genes with a conserved C-terminal EH-i-like motif. Similarly, *foxQ2-1* in *C. teleta* and *foxQ2-11* in *O. fusiformis* are phylogenetically related and both show maternal expression. Therefore, these findings support that some of the expansions of *foxQ2* genes are shared across spiralian lineages and that the expansion of this class of *Fox* genes may also resulted in the evolution of novel expression dynamics.

**Fig. 5.**
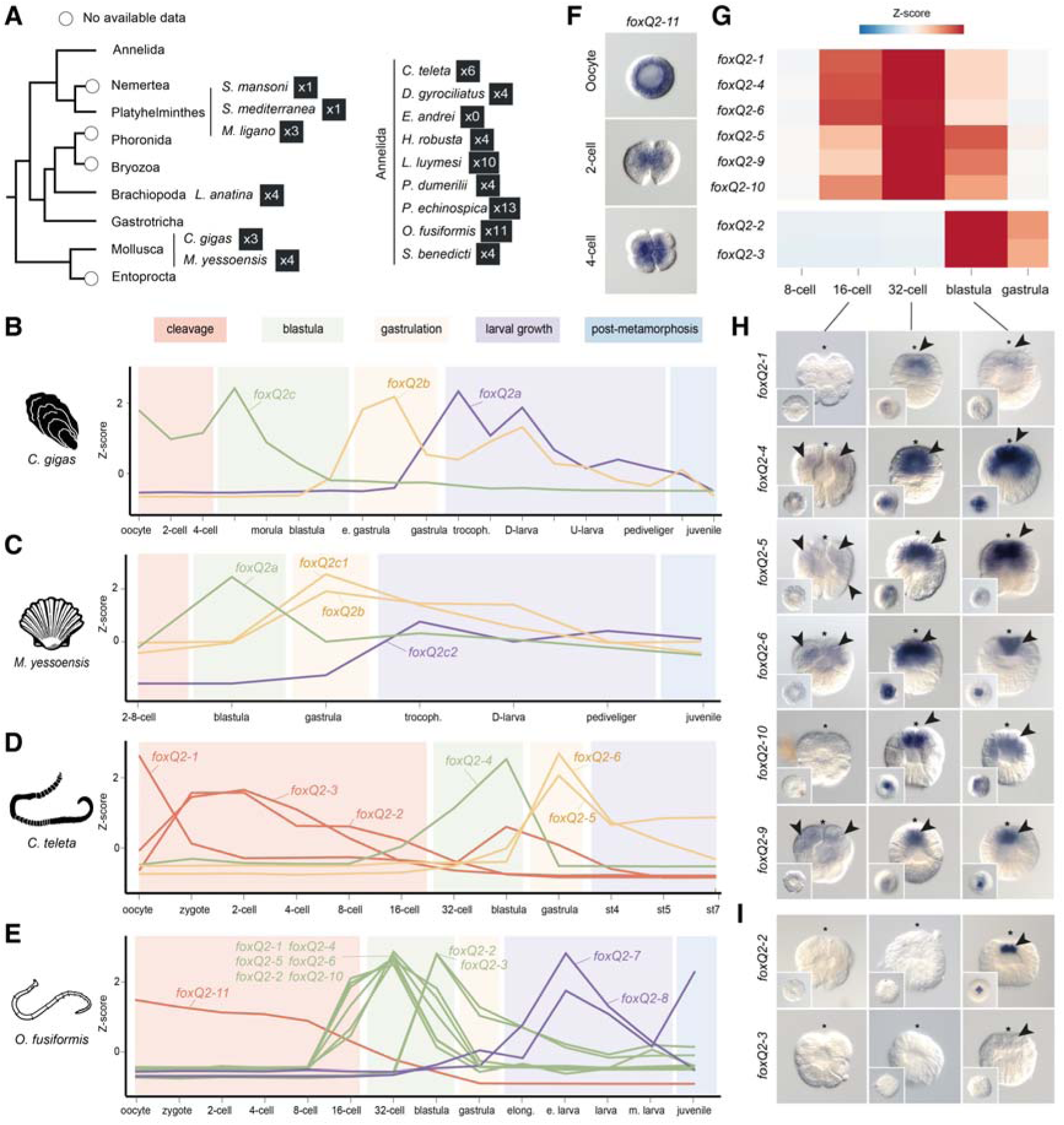
The *foxQ2* gene complement in Annelida and Mollusca. (**A**) Number of *foxQ2* genes in selected spiralian taxa based on the orthology assignment in Figure 4A, under a recent phylogenetic framework (Marlétaz et al, 2019). **(B**–**E)** Line plots representing z-score values of the temporal expression dynamics of *foxQ2* paralogs during the development of the molluscs *C. gigas* and *M. yessoensis*, and the annelids *C. teleta* and *O. fusiformis*. Exact times of development are detailed in Supplementary Table 3. Major developmental phases are indicated in coloured boxes to help compare phases of development across the four species (early cleavage in pink, blastula in green, gastrulation in yellow, larval growth in purple and post-metamorphosis in blue). The line of expression of each gene is coloured according to the stage in which it shows a peak of expression. (**F**) Apical views of whole mount *in situ* hybridisation of *foxq2-11* in *O. fusiformis* at the oocyte stage, the 2-cell stage, and the 4-cell stage. This gene shows maternal expression and equal distribution in early blastomeres. (**G**) Heatmap of expression of *foxQ2* paralogs with peaks of expression at 4 and 5 hours post fertilisation (blastula) in *O. fusiformis*. Colours show the normalized z-score value of expression, with red blocks indicating high levels of expression and blue blocks indicating low levels of expression. (**H**, **I**) Lateral views of whole mount *in situ* hybridisation of *foxQ2* paralogs expressed from 3 to 5 hours post fertilisation (blastula). Insets show apical views, the asterisks point to the animal/apical pole, and arrowheads to the domains of expression.

We next analyse whether *foxQ2* paralogs also showed varying spatial expression domains in *O. fusiformis* using whole mount *in situ* hybridization across targeted developmental stages. Except for *foxQ2-8* and *foxQ2-7*, for which we did not observe any clear expression pattern at the larval stage, all other nine *foxQ2* paralogs showed distinct domains of expression. The paralog *foxQ2-11*, which has a high expression in the oocyte, is detected in the zygote and appears equally distributed in all blastomeres at the 2-cell and 4- cell stages (Figure 5F). The genes *foxQ2-1, foxQ2-4, foxQ2-5*, *foxQ2-6*, *foxQ2-9*, *foxQ2-10*, which peak at 4 hours post fertilisation (Figure 5G) and are probably independent expansions in *O. fusiformis* (Figure 4A), are expressed at the apical pole, but encompassing varying areas of expression, from ones restricted to just the animal micromeres (e.g., *foxQ2-6* and *foxQ2-9*) to nearly the entire animal hemisphere (e.g., *foxQ2-4*, *foxQ2-5* and *foxQ2-6*) (Figure 5H). Finally, the expression of the two *foxQ2* paralogs with a C-terminal EH-i-like motif—*foxQ2-2* and *foxQ2-3*—is restricted to the apical-most micromeres from 5 hours post fertilisation onwards, consistent with the observed temporal dynamics and its regulation by ERK1/2 activity at that timepoint (Figure 5G, I). Altogether, our gene expression analyses support that the multiple copies of *foxQ2* have retained the evolutionarily conserved expression of this *Fox* gene class in anterior/apical development (Marlow et al. 2014), yet they have probably undergone temporal, spatial, and regulative sub-functionalisation.

## Discussion

### The evolution of the *Fox* gene complement in Annelida and Spiralia

Taking advantage of the reference genome assembly and comprehensive functional data available for the annelid *O. fusiformis* (Liang et al. 2022), our study identified and characterised the full *Fox* gene repertoire in a member of Oweniidae, the sister lineage to all remaining annelids (Rouse et al. 2022). The 34 *Fox* genes of *O. fusiformis*, belonging to 21 of the 23 classes predicted for the bilaterian ancestor (Kaestner et al. 2000; Benayoun et al. 2011), are the most complete *Fox* gene repertoire for a spiralian reported to date, and thus helps to clarify the ancestral *Fox* gene complement in Spiralia and Lophotrochozoa (Marlétaz et al. 2019), as well as the evolution of certain *Fox* gene classes in this animal group. All 23 *Fox* gene classes were present in the last common Spiralian ancestor, including a putative divergent class in some previous reports referred to as *foxY* (Wu et al. 2020; Mei, Fei, Jun, Li, Yang, Xu, Liu, Que, Li, Zhang, et al. 2014). Given the low node support for this clade in phylogenetic analyses, putative *foxY* genes likely represent fast evolving lineage-specific *Fox* genes, instead of a *bona fide* class. Alternatively, *foxY* genes might have a common evolutionary origin—blurred by the fast rate of molecular evolution of the members in this class—and share developmental roles (e.g., *foxY* in both echinoderms, molluscs and *O. fusiformis* are expressed maternally) (Song & Wessel 2012; Mei, Fei, Jun, Li, Yang, Xu, Liu, Que, Li & Zhang 2014). Either way, further studies of *foxY* genes are needed to help to clarify these scenarios, as well as the expression and role of this divergent group of Forkhead containing genes.

Our study supports that lineage-specific losses and expansions are common in the evolution of the *Fox* genes in Spiralia. While a small number of *Fox* gene classes have been independently lost in some annelid lineages (e.g., *foxE* and *foxI* in *O. fusiformis* and *foxN1/4*, *foxO* and *foxP* in *C. teleta*) (Mei, Fei, Jun, Li, Yang, Xu, Liu, Que, Li, Zhang, et al. 2014), Mollusca (or at least Conchifera) has experienced ancestral losses of two classes, namely *foxI* and *foxQ1*, as well as independent lineage-restricted losses (e.g., *foxG* and *foxM* in *C. gigas*, *foxO* in *L. gigantea*) (Mei, Fei, Jun, Li, Yang, Xu, Liu, Que, Li, Zhang, et al. 2014; Shimeld, Boyle, et al. 2010). Consistent with its faster rate of molecular evolution, the platyhelminth *S. mediterranea* has a more degraded *Fox* gene repertoire, with up to nine *Fox* classes missing (*foxAB*, *foxB*, *foxE*, *foxH*, *foxL2*, *foxQ1*, *foxJ2/3*, *foxM* and *foxN1/4*) (Pascual-Carreras et al. 2021). Compared to the annelid *C. teleta* and molluscs, *O. fusiformis*, however, has experienced more duplications of certain *Fox* classes (*foxAB*, *foxN1/4* and *foxN2/3*), a phenomenon that is more common in the platyhelminth *S. mediterranea* (Pascual-Carreras et al. 2021). Notably, and as discussed below, expansions of the *foxQ2* class are common in Annelida and Mollusca, with some lineages, including *O. fusiformis*, exhibiting large lineage-specific expansions. Therefore, while Annelida and Mollusca tend to display a relatively stable complement of *Fox* genes, other spiralian lineages are more divergent and dynamic. Future work on the expression and function of *Fox* genes across spiralian lineages will help to clarify how these expansions and losses impact developmental programmes and the diversification of body plans in Spiralia.

Gene architecture differences between clade I and II and the ancestral chromosomal linkage of the *foxC*, *foxF*, *foxL1* and *foxQ1* classes characterise the evolution of *Fox* genes (Larroux et al. 2008; Shimeld, Boyle, et al. 2010). While our gene architecture data generally support the previous observations that the clade I of *Fox* genes are intronless, *O. fusiformis*— differently from the annelid *C. teleta*—does not exhibit a *Fox* gene cluster involving *foxC*, *foxF*, *foxL1* and *foxQ1* (Shimeld, Boyle, et al. 2010). This contrasts with the overall conservation of gene macrosynteny in *O. fusiformis* (Liang et al. 2022) and suggests that some intra-chromosomal rearrangements might have happened in this species. Despite the lack of a cluster organisation, however, these genes retain a mesodermal expression in *O. fusiformis* (at least for *foxC*, *foxF*, *foxL1*; see below), as observed in *C. teleta* and other spiralians (Shimeld, Boyle, et al. 2010). Therefore, our study challenges the hypotheses that the concerted mesodermal co-expression of *foxC*, *foxF*, *foxL1* and *foxQ1* genes was the selective pressure for maintaining their cluster integrity (Shimeld, Boyle, et al. 2010; Shimeld, Degnan, et al. 2010), suggesting instead that more local, gene-specific regulation is responsible for their expression dynamics.

### The expression dynamics of the *Fox* gene complement in *O. fusiformis*

By combining new and previously reported temporal and spatial expression data in *O. fusiformis*, our study provides a comprehensive view of the developmental dynamics of *Fox* genes in this annelid species, suggesting conserved and potentially new developmental roles for certain *Fox* classes. In most animals studied to date, including annelids, molluscs, brachiopods, phoronid, bryozoans, planarians and nemertean species, *foxA* is a key effector of foregut formation and a marker of endodermal tissues (Jr Kai Yu et al. 2008; Adler et al. 2014; Fritzenwanker et al. 2014; Boyle & Seaver 2008, 2010; Arenas-Mena 2006; Kostyuchenko et al. 2019; Kwak et al. 2018; Lartillot et al. 2002; Martín-Durán et al. 2015, 2016; Perry et al. 2015; Andrikou et al. 2019; Vellutini et al. 2017; Fuchs et al. 2011; Pascual-Carreras et al. 2021). In *O. fusiformis*, *foxA* is first detected in the vegetal macromeres at the blastula stage and later observed in the gastrula endoderm (peak of expression) and in the mouth and midgut of the developing larvae (Martín-Durán et al. 2016), thus supporting the role of the *foxA* class in endoderm and gut formation in animals. Similarly, the gene *foxJ1* is involved in ciliogenesis in a number of animals (Choksi et al. 2014; Xianwen Yu et al. 2008), including during the formation of the prototroch in the annelid *P. dumerilii* (Marlow et al. 2014). In *O. fusiformis*, *foxJ1* is expressed soon after gastrulation in the presumptive prototroch precursors and later on in the heavily ciliated cell types of the larva (the apical organ and prototroch) (Carrillo-Baltodano et al. 2021), reinforcing a conserved role of this *Fox* gene in the development of ciliated organs in Annelida and metazoans in general.

Existing gene expression data also support a conserved role of several *Fox* gene classes in mesoderm development in *O. fusiformis*. The *foxH* class regulates mesoderm development and embryonic organising activity in vertebrates (Hoodless et al. 2001; Pogoda et al. 2000; Kofron et al. 2004), and it is downstream of the embryonic organiser and likely involved in mesoderm and posterodorsal development in *O. fusiformis* (Seudre et al. 2022), with a similar temporal expression dynamic reported in the oyster *C. gigas* (Mei, Fei, Jun, Li, Yang, Xu, Liu, Que, Li & Zhang 2014). Similarly, the temporal and spatial expression domains of *foxL1*, *foxC* and *foxF* support their role in mesoderm formation in *O. fusiformis* (Martín-Durán et al. 2016), as described in other spiralians and metazoans (Shimeld, Boyle, et al. 2010; Wotton et al. 2008; Wotton & Shimeld 2011; Fritzenwanker et al. 2014), albeit they do not retain their linked chromosomal position in this annelid species. Notably, *foxQ1* which is absent from all molluscan species studied to date, is expressed in the stomodeum and pharynx in *C. teleta* (Shimeld, Boyle, et al. 2010; Wu et al. 2020; Mei, Fei, Jun, Li, Yang, Xu, Liu, Que, Li, Zhang, et al. 2014). However, *foxQ1* is only expressed at the juvenile stage in *O. fusiformis*, suggesting that the role of this *Fox* gene class might differ between annelid and spiralian species.

The expression of *Fox* genes in other annelids and spiralians combined with the temporal dynamics of orthologs in *O. fusiformis* provide evidence of the potential roles of certain *Fox* gene classes in this annelid species. For instance, *foxD* is involved in myogenesis and ventral patterning in the annelid *P. dumerilii*, the platyhelminth *S. mediterranea* and the brachiopod *T. transversa* (Passamaneck et al. 2015; Pascual-Carreras et al. 2021; Lauri et al. 2014) and it becomes expressed after gastrulation and specially during organogenesis and initiation of myogenesis in *O. fusiformis* (Carrillo-Baltodano et al. 2021). The *foxO* class is a regulator of cell death in the planarian *S. mediterranea* (Pascual-Carreras et al. 2021) and a effector of cell division during early cleavage in the annelid *H. austinensis* (Kwak et al. 2018). In *O. fusiformis*, *foxO* is also expressed maternally and during early cleavage, as well as during embryonic periods with active cell turnover (Carrillo-Baltodano et al. 2021). The expression of the *foxAB* class has been only studied in the annelid *C. teleta*, which has a single ortholog that becomes expressed in a unique D-quadrant cell during early cleavage and is later involved in ectoderm differentiation and foregut formation (Boyle et al. 2014). *Owenia fusiformis* has instead two *foxAB* paralogs, one (*foxAB-a*) exclusively expressed at the time of the specification of the organiser cell at 5 hours post fertilisation and a second one (*foxAB-b*) that peaks at the gastrula stage before gradually fading away during larval development. Although further expression analyses are needed, we speculate that the two paralogs in *O. fusiformis* could play similar roles than those described in *C. teleta*, with *foxAB-a* acting during the axial body patterning and *foxAB-b* being involved in ectoderm and foregut formation later in embryogenesis.

Our comprehensive developmental time course of the *Fox* genes in *O. fusiformis* also uncovered dynamics of expression for *Fox* gene classes for which there is little understanding of their roles during annelid and spiralian embryogenesis. For instance, *foxJ3*, *foxM1*, *foxN2/3-b* and *foxN1/4-a* and *foxN1/4-b* show maternal expression, which is maintained until the 8-cell stage, suggesting that they might be potential regulators of early cleavage and/or cell fate specification in this species. The genes *foxG1* and *foxP* have a narrow peak of expression at the time of the specification of the embryonic organiser in *O. fusiformis* (Seudre et al. 2022), and could therefore be involved in the establishment of the embryonic polarity and body plan in this annelid. Finally, some *Fox* genes are restricted to either the larva (*foxK2*) or the juvenile (*foxB, foxL2* and *foxQ1*), yet their potential role at those stages remains unknown. Altogether, our study sets the stage for further expression and functional studies of *Fox* genes in *O. fusiformis* and spiralian embryogenesis, which ultimately will help to better understand the plasticity and development roles of this major family of transcription factors in animal development and evolution.

### The complex evolutionary history of *foxQ2* genes

The *foxQ2* class often comprises at least two paralogs in many cnidarian and bilaterian lineages studied to date, except in Tetrapoda, which lost this *Fox* gene class (Fritzenwanker et al. 2014; Santagata et al. 2012; Yu et al. 2003; Kitzmann et al. 2017; Sinigaglia et al. 2013; Marlow et al. 2014; Chevalier et al. 2006; Mazet et al. 2003). Notably, *foxQ2* genes show a remarkable conservation of their expression patterns across phylogenetically distant animal lineages. In deuterostomes like the cephalochordate *Branchiostoma floridae*, the echinoderm *Strongylocentrotus purpuratus* and the hemichordate *Saccoglossus kowalevskii, foxQ2* genes are expressed in the apical pole during embryogenesis (Range & Wei 2016; Fritzenwanker et al. 2014), and *foxQ2* genes also play central roles in anterior neuronal development in the insects *Drosophila melanogaster* and *Tribolium castaneum*, and the spider *Parasteatoda tepidariorum* (Kitzmann et al. 2017; Schacht et al. 2020; Lee & Frasch 2004). In the cnidarians *Nematostella vectensis* and *Clytia hemispherica*, *foxQ2* genes are expressed in and required for the proper development of the aboral pole, in support for the—still debated—homology between the cnidarian aboral pole and the bilaterian anterior pole (Chevalier et al. 2006; Sinigaglia et al. 2013). In Spiralia, *foxQ2* genes share an apical expression in the annelid *P. dumerilii* and the brachiopod *T. transversa* (Santagata et al. 2012; Marlow et al. 2014), which, as our study shows, is consistent with the majority of expression domains for *foxQ2* genes observed in the annelid *O. fusiformis*. Therefore, *foxQ2* genes appear to participate in ancient and broadly conserved genetic programmes for apical and axial patterning in metazoans.

The consistent expression of *foxQ2* genes in apical territories contrasts, however, with the complex phylogenetic pattern of evolution of this class of *Fox* genes. As our study reveals, expansions of the *foxQ2* class are common in Spiralia, and specially in Annelida, with 11 paralogs in *O. fusiformis*, and 10 and 13 in the vestimentiferans *L. luymesi* and *P. echinospica*, respectively. While many of these paralogs probably emerged from species-specific expansions (e.g., *foxQ2-1*, *foxQ2-4*, *foxQ2-5*, *foxQ2-6*, and *foxQ2-9* in *O. fusiformis*; and the cluster of vestimentiferan *foxQ2* paralogs with an N-terminal EH-i-like motif) other paralogs might have a more ancient origin, tracing back to Annelida (e.g., *foxQ2-1* in *C. teleta* and *foxQ2-11* in *O. fusiformis*, both expressed maternally) and even Spiralia (e.g., *foxQ2-2* in *C. teleta* and *foxQ2-c2* in *M. yessoensis*). Notably, however, nearly all bilaterian and cnidarian lineages retain at least a copy (two in the annelids *O. fusiformis* and *D. gyrociliatus*) of a subclass of *foxQ2* genes with a C-terminal EH-i-like motif. In *O. fusiformis,* these two genes (*foxQ2-2* and *foxQ2-3*) are controlled by the ERK1/2 signalling that establishes the axial polarity of the embryo and consequently show a narrow peak of expression at the animal pole at the time of the specification of the organizer cell at five hours post fertilisation (Seudre et al. 2022). Based on these observations, we propose a model in which an ancestral *foxQ2* gene containing a C-terminal EH-i-like motif originated in the last common ancestor to Cnidaria and Bilateria, followed by independent expansions in certain bilaterian and cnidarian lineages and fast divergence of the new copies, which tended to lose the EH-i-like motif. Despite these duplications, however, the new paralogs did not generally acquire radically different functions (i.e., neofunctionalization) but rather retain a role in aboral/apical/anterior development, and evolved temporal, spatial and regulative specialisation of their expression (i.e., subfunctionalisation), as observed for example in *O. fusiformis* and the hemichordate *S. kowalevskii* (Fritzenwanker et al. 2014). Further analyses of the gene regulatory network associated with *foxQ2* genes in a broader range of metazoans, especially in spiralians with multiple copies, and role of the EH-i-like motif will contribute to uncover the evolutionary history and developmental consequences of *foxQ2* expansions during animal diversification.

Altogether, our study informs the evolution, temporal, and spatial expression of the largely conserved *Fox* gene repertoire in the oweniid annelid *O. fusiformis*. Our findings provide valuable information to reconstruct the ancestral complement of these core developmental regulators in Spiralia and to continue unravelling the embryological role and contribution of this major family of transcription factors to the evolution of animal body plans.

## Material and Methods

### Animal husbandry and embryo collection

Adult specimens of *Owenia fusiformis* Delle Chiaje, 1844 were collected and shipped to London from the coast near the Station Biologique de Roscoff (France) during their reproductive season (May to July). In the lab, animals were kept in aquaria with mud and artificial seawater (ASW) at 15°C. *In vitro* fertilizations were conducted as previously described (Martín-Durán et al. 2016; Carrillo-Baltodano et al. 2021) and embryos were kept in glass bowls at 19°C until they reach the desired developmental stage. Larval stages were relaxed in 8% MgCl2 and all embryonic samples fixed in 4% formaldehyde in sea water (or MgCl2, for larvae) for 1 hour at room temperature. After washing the fixative with phosphate buffer saline (PBS) supplemented with 0.1% Tween-20, embryos and larvae were dehydrated to 100% methanol and stored at -20°C.

### Identification of *forkhead* genes and orthology assignment

Candidate *Fox* genes for *O. fusiformis* were initially retrieved from the functional annotation of its genome assembly (European Nucleotide Archive, accession number: GSE184126) (Liang et al. 2022), and the *foxC* sequence was obtained from a previous study (Martín-Durán et al. 2016). The annotated *Fox* sequences from spiralians (*Capitella teleta*, *Crassostrea gigas*, *Lottia gigantea*, *Mizuhopecten yessoensis*, *Terebratallia transversa, Crepidula fornicata, Lingula unguis*), ecdysozoans (*Drosophila melanogaster, Strigamia maritima, Caenorhabditis elegans, Tribolium castaneum)*, deuterostomes (*Homo sapiens*, *Danio rerio, Mus musulus, Branchiostoma lanceolatum, Saccoglossus kowalevskii, Strongylocentrotus purpuratus*) and the cnidarian *Nematostella vectensis* were identified by mining published transcriptomes and databases (Supplementary Figures 7, 8). All the above Fox proteins were searched against the Pfam database to trim the Forkhead domain (PF00250). Multiple protein alignments (Supplementary Figures 7, 8) were performed with MAFFT v.7 (Katoh & Standley 2013) with the L-INS-i strategy and computed independently for *Fox* genes belonging to the Clade I and Clade II. Poorly aligned regions were removed with gBlocks version 0.91b (Talavera et al. 2007) and maximum likelihood trees were constructed with RAxML v.8.2.11 (Stamatakis 2014) with automatic identification of the model of protein evolution and automatic bootstrapping. Both alignments were modelled with an LG matrix (Le & Gascuel 2008). Bayesian reconstructions in MrBayes v.3.2.7a (Ronquist & Huelsenbeck 2003) were also performed using the same LG matrix as a prior. For each clade, two runs with four chains (one cold, three hot) were run for 70,000,000 generations. Resulting trees were visualized and edited with FigTree (https://github.com/rambaut/figtree/).

### Genomic structure of *Fox* genes

The exon and intron positions, as well as the chromosomal location for each *Fox* gene were determined based on the gene annotation of the genome assembly of *O. fusiformis* (Liang et al. 2022) using the Integrative Genomics Viewer (IGV) (Robinson et al. 2011). The position of the Forkhead domain within each *Fox* gene was determined using the conserved domain database CDD/SPARCLE (Lu et al. 2020). The genomic architecture of the *Fox* genes was visualized using the online software IBS (Liu et al. 2015) and transferred to Illustrator Creative Cloud 2022 (Adobe).

### Phylogenetic analysis of the *foxQ2* family

To study the evolution of the FoxQ2 family in Spiralia, we retrieved sequences from spiralians (C. teleta, C. gigas, Dimorphilus gyrociliatus, Eisenia andrei, Helobdella robusta, L. gigantea, L. luymesi, M. yessoensis, Platynereis dumerilii, P. echinospica, Streblospio benedicti), ecdysozoans (D. melanogaster, N. vetripennis, P. humanus, T. castaneum), deuterostomes (O. latipes, S. kowalevskii, S. purpuratus, B. floridae) and cnidarians (C. hemisphaerica, H. vulgaris, N. vectensis) from published genomes, transcriptomes and databases. For all genes, membership to the foxQ2 family was manually confirmed using the conserved domain database CDD/SPARCLE (Lu et al. 2020) and identifying the presence of a full FoxQ2 specific domain (Accession number CD20035) within the sequences. Some genes previously misannotated and assigned to other Fox gene classes were renamed as foxQ2. To identify the EH-I like domain in the foxQ2 genes, we generated a EH-1-like motif position-specific scoring matrix (PSSM) using STREME v 5.4.1 (Bailey 2021), with a motif width of 7 to 10 amino acid and retaining motifs with p-value < 0.05, and confirmed the identified motif by comparison with results obtained by (Yaklichkin et al. 2007) (Figure 4B). The position and the presence of the EH-like motifs within the FoxQ2 sequences were determined by inputting the EH-i-like domain matrix into FIMO v 5.4.1 (Grant et al. 2011), matching motifs with a q-value < 0.05. For phylogenetic analyses, multiple protein alignments were performed with MAFFT v.7 as explained above and trees were reconstructed from a set of sequences selected with a Q.insect amino acid replacement matrix (Minh et al. 2021) to account for transition rates, the gamma distribution with four categories (G4) (Yang 1994) to describe sites evolution rates, and an optimisation of amino acid frequencies using maximum likelihood in IQ-TREE v.2.1.2 (Minh et al. 2020). A thousand ultrafast bootstraps were used to extract branch support values and posterior probabilities were obtained through Bayesian reconstruction in MrBayes v.3.2.7a (Ronquist & Huelsenbeck 2003) as described above using the general time reversible (GTR) model as a prior and 50,000,000 generations.

### Gene expression developmental time course

Stage-specific RNA-seq data covering 14 time-points, from the unfertilized oocyte to the juvenile stage (Liang et al. 2022) were used to retrieve gene expression dynamics for all *Fox* genes in *O. fusiformis*. The transcriptomes of multiple developmental stages of the molluscs *C. gigas* and *M. yessoensis* where retrieved from Gene Expression Omnibus (accession number GSE31012) (Wang et al. 2012) and the Short Read Archive database (accession numbers SRX1026991, SRX2238787 to SRX2238809, SRX2250256 to SRX2250259, SRX2251047, SRX2251049, SRX2251056, SRX2251057 and SRX2279546) (Wang et al. 2017), respectively (Supplementary Table 3). The developmental expression profiles of *foxQ2* paralogs in *C. teleta* were computed using stage-specific RNA-seq data (Liang et al. 2022) (Supplementary Table 4). For all four species, if more than one sample were collected for a given developmental stage, the values of expression of the different replicates were averaged. The timing of sample collection and the number of replicates for each species and stages is detailed in Supplementary Table 3. Heatmaps were generated using the package pheatmap v.1.0.12 available in R, where colour intensity shows the z-score value for each candidate genes (blue: low expression, red: high expression) (Kolde, 2015).

### Gene isolation and whole mount *in situ* hybridization

*foxQ2* genes in *O. fusiformis* were amplified using gene specific primers, producing DNA templates for riboprobe synthesis by successive rounds of nested PCR on cDNA obtained from mixed developmental stages as initial template. Riboprobes were synthesized with the T7 enzyme following manufacturer’s recommendations (Ambion’s MEGAscript kit, #AM1334) and stored in hybridization buffer at a concentration of 50 ng.μl-1 at -20°C. Single colorimetric *in situ* hybridization of embryos and mitraria larvae were performed following an established protocol (Martín-Durán et al. 2016; Carrillo-Baltodano et al. 2021; Seudre et al. 2022).

### Imaging

Representative embryos from colorimetric whole mount *in situ* hybridization were cleared in 70% glycerol in PBS and imaged with a Leica DMRA2 upright epifluorescent microscope equipped with an Infinity5 camera (Lumenera), using bright field, differential interference contrast (DIC) optics. Brightness/contrast and colour balance were adjusted using Pixelmator Pro (v. 2.0.3) and applied to the whole image, not parts. Final figure panels were designed using Illustrator Creative Cloud 2022 (Adobe).

## Data availability

The data underlaying this article are available in the article and in its online supplementary material.

## Author contributions

OS and JMM-D designed the study; OS contributed to all analyses and drafted the manuscript; FMM-Z contributed to phylogenetic analyses; VR and AMC-B helped with gene expression analyses; JMM-D contributed to data analyses and manuscript writing; all authors contributed to data interpretation and manuscript writing.

## Acknowledgements

We thank all members of the Martín-Durán lab for their support, and in particular Yan Liang for her help with mapping gene expression data in the genome annotation of *O. fusiformis*. We also thank the Station Biologique de Roscoff for their help with collections and animal supplies. This work was funded by the Horizon 2020 framework programme to JMM-D (European Research Council Starting Grant action number 801669). In addition, the research leading to these results received funding for the European Union’s Horizon 2020 research and innovation programme under grant agreement No 730984, ASSEMBLE Plus project.

## Supplementary Figure Legends

**Supplementary Fig. 1 – Maximum likelihood orthology analysis of *O. fusiformis* Clade I *Fox* genes.** Maximum likelihood phylogenetic analysis for Clade I *Fox* genes members in *O. fusiformis* and 18 other metazoan species, using the *foxK* class as outgroup. Coloured boxes show monophyletic clades corresponding to previously described *Fox* classes. *Owenia fusiformis* sequences are highlighted in red. Note that the *foxQ2* class is expanded in this species. Only bootstrap values at nodes that supposedly represent distinct orthology groups are shown. High support values within orthogroups (e.g., between paralogs) were omitted for the sake of clarity. Scale bar indicates the number of amino acid substitutions per site alongside the branches.

**Supplementary Fig. 2 – Maximum likelihood orthology analysis of *O. fusiformis* Clade II *Fox* genes.** Maximum likelihood phylogenetic analysis for Clade II *Fox* genes members in *O. fusiformis* and 18 other metazoan species, using as outgroup the *foxG* class. Coloured boxes show monophyletic clades corresponding to previously described *Fox* classes. *Owenia fusiformis* sequences are highlighted in red. Only bootstrap values at nodes that supposedly represent distinct orthology groups are shown. High support values within orthogroups (e.g., between paralogs) were omitted for the sake of clarity. Scale bar indicates the number of amino acid substitutions per site alongside the branches.

**Supplementary Fig. 3 – Bayesian orthology assignment of *O. fusiformis* Clade I *Fox* genes.** Bayesian phylogenetic analysis for Clade I *Fox* genes members as in Supplumentary Figure 1. Coloured boxes show monophyletic clades corresponding to previously described *Fox* classes. *Owenia fusiformis* sequences are highlighted in red. Only posterior probability values at nodes that supposedly represent distinct orthology groups are shown. High posterior probability values within orthogroups (e.g., between paralogs) were omitted for the sake of clarity. Scale bar indicates the number of amino acid substitutions per site alongside the branches.

**Supplementary Fig. 4 – Bayesian orthology assignment of *O. fusiformis* Clade I *Fox* genes.** Bayesian phylogenetic analysis for Clade I *Fox* genes members as in Supplementary Figure 2. Coloured boxes show monophyletic clades corresponding to previously described *Fox* classes. *Owenia fusiformis* sequences are highlighted in red. Only posterior probability values at nodes that supposedly represent distinct orthology groups are shown. High posterior probability values within orthogroups (e.g., between paralogs) were omitted for the sake of clarity. Scale bar indicates the number of amino acid substitutions per site alongside the branches.

**Supplementary Fig. 5 – Maximum likelihood phylogenetic analysis of *foxQ2* genes.** Maximum likelihood analysis for *foxQ2* genes containing an intact FoxQ2 domain (accession number CD20035) across metazoan lineages, using the *foxL2* family as the outgroup, as in Figure 4. Sequence names are coloured based on the position of the EH-i-like motif (shown in Figure 4B) with respect to the FoxQ2 box (sequences with a C-terminal position of the EH-I-like motif are in gray; sequences with an N-terminal position of the EH-I-like motif are in green; sequences lacking the EH-i-like motif are in black). The gray box highlights the monophyletic clade largely containing sequences of both cnidarian and bilaterian lineages with a C-terminal EH-i-like motif. Red dots indicate *O. fusiformis foxQ2* paralogs. Only bootstrap values at nodes that supposedly represent distinct orthology groups are shown. High support values within orthogroups (e.g., between paralogs) were omitted for the sake of clarity. Scale bar indicates the number of amino acid substitutions per site alongside the branches.

**Supplementary Fig. 6 – Bayesian phylogenetic analysis of *foxQ2* genes.** Bayesian reconstruction of the evolution of *foxQ2* genes containing an intact FoxQ2 domain (accession number CD20035) across metazoan lineages, using the *foxL2* family as the outgroup, as in Figure 4. Sequence names are coloured based on the position of the EH-i-like motif (shown in Figure 4B) with respect to the FoxQ2 box (sequences with a C-terminal position of the EH-I-like motif are in gray; sequences with an N-terminal position of the EH-I-like motif are in green; sequences lacking the EH-i-like motif are in black). The gray box highlights the monophyletic clade largely containing sequences of both cnidarian and bilaterian lineages with a C-terminal EH-i-like motif. Red dots indicate *O. fusiformis foxQ2* paralogs. Only posterior probability values at nodes that supposedly represent distinct orthology groups are shown. High posterior probability values within orthogroups (e.g., between paralogs) were omitted for the sake of clarity. Scale bar indicates the number of amino acid substitutions per site alongside the branches.

**Supplementary Fig. 7 – Multiple Protein Alignment for Clade I *Fox* genes.** Aligned sequences were trimmed and processed in gBlocks to render this final alignment. Residues are coloured following the ClustalX colour scheme. Bar plots of residue conservation, occupancy, quality, together with the frequency of the consensus amino acid for each position are depicted below the alignment. The symbol “+” denotes a fully conserved residue.

**Supplementary Fig. 8 – Multiple Protein Alignment for Clade II *Fox* genes.** Aligned sequences were trimmed and processed in gBlocks to render this final alignment. Residues are coloured following the ClustalX colour scheme. Bar plots of residue conservation, occupancy, quality, together with the frequency of the consensus amino acid for each position are depicted below the alignment. The symbol “+” denotes a fully conserved residue.

**Supplementary Fig. 9 – Multiple Protein Alignment for *foxQ2* genes.** Aligned sequences were trimmed to the FoxQ2 domain to render this final alignment. Residues are coloured following the ClustalX colour scheme. Bar plots of residue conservation, occupancy, quality, together with the frequency of the consensus amino acid for each position are depicted below the alignment. The symbol “+” denotes a fully conserved residue.

## Notes

### Competing Interest Statement

The authors have declared no competing interest.

