## Supplementary figures and images for "The *Fox* gene repertoire in the annelid *Owenia fusiformis* reveals multiple expansions of the *foxQ2* class in Spiralia"

### Supplementary Figures 1 to 9

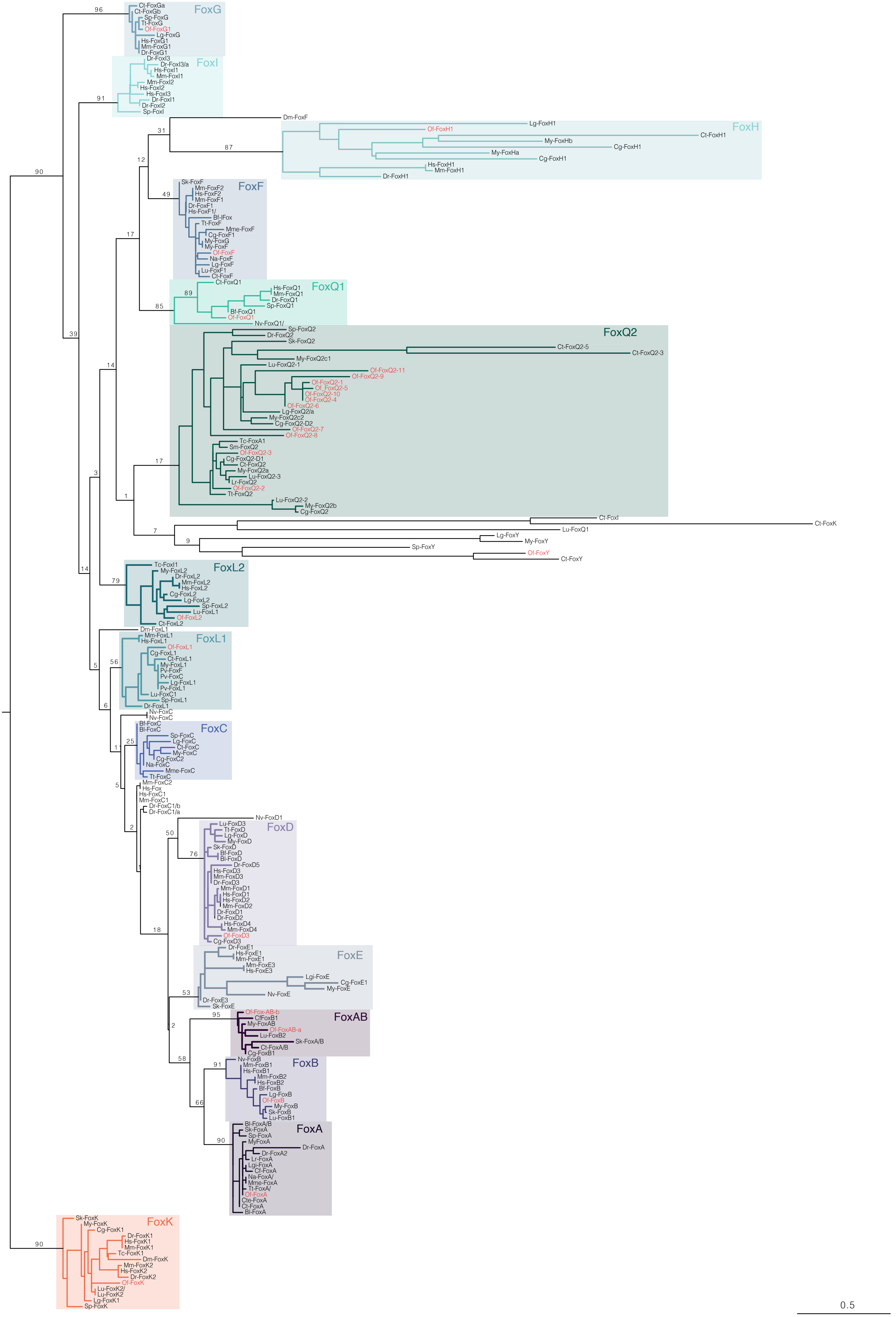

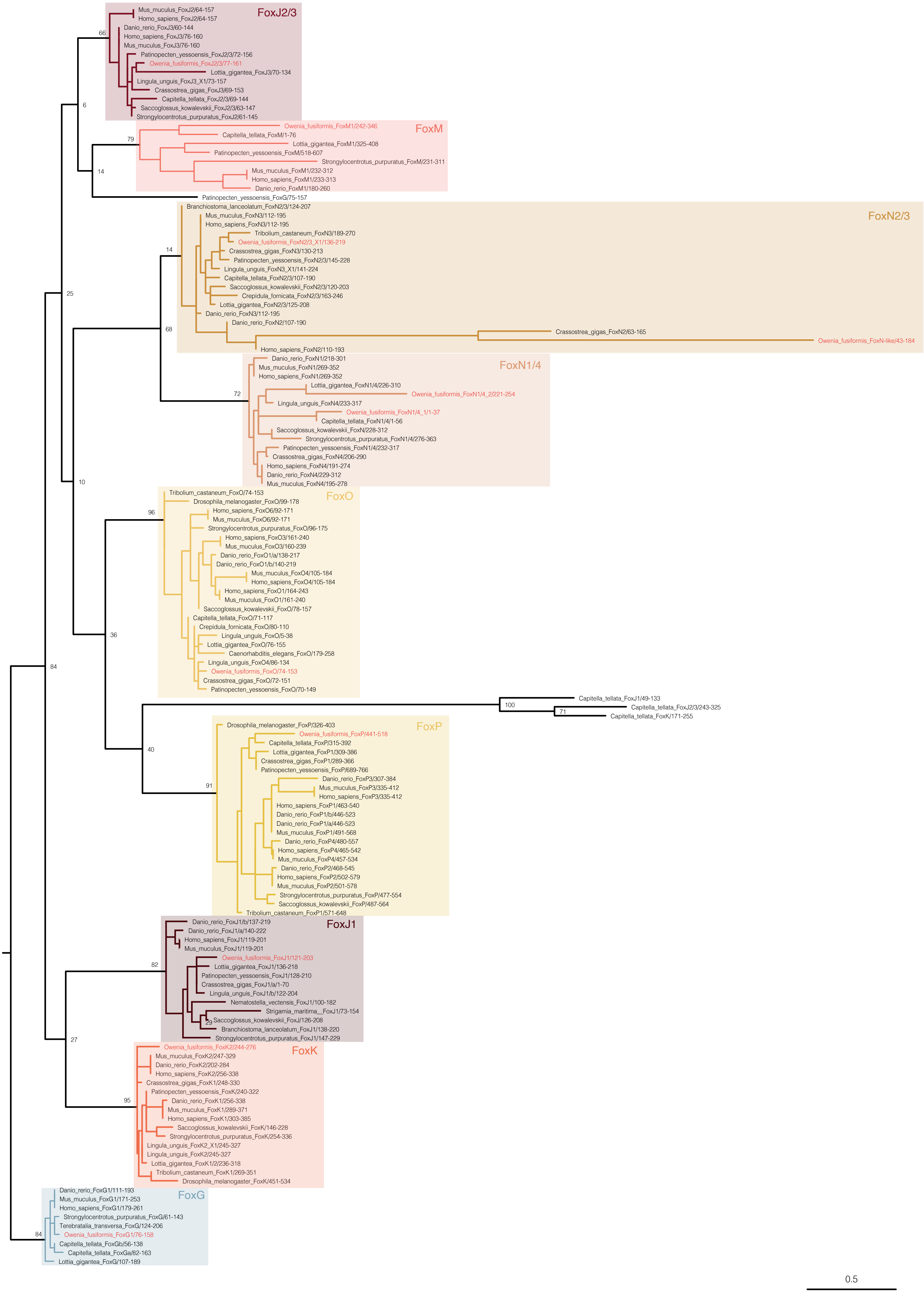

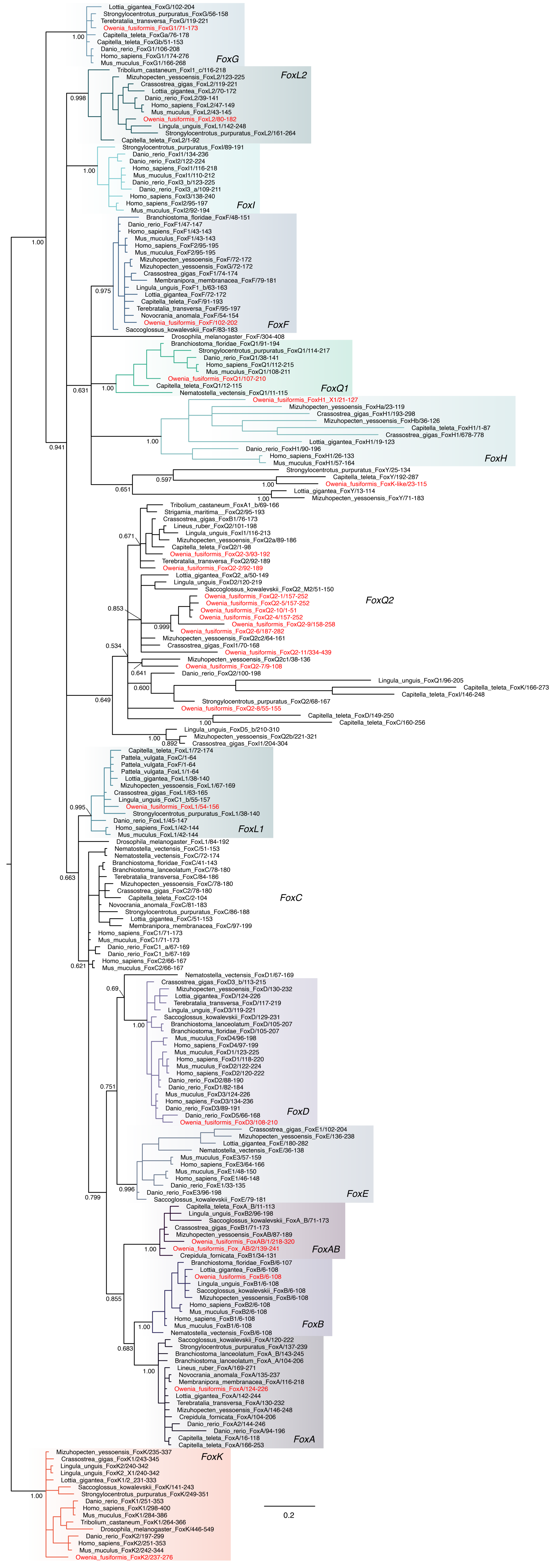

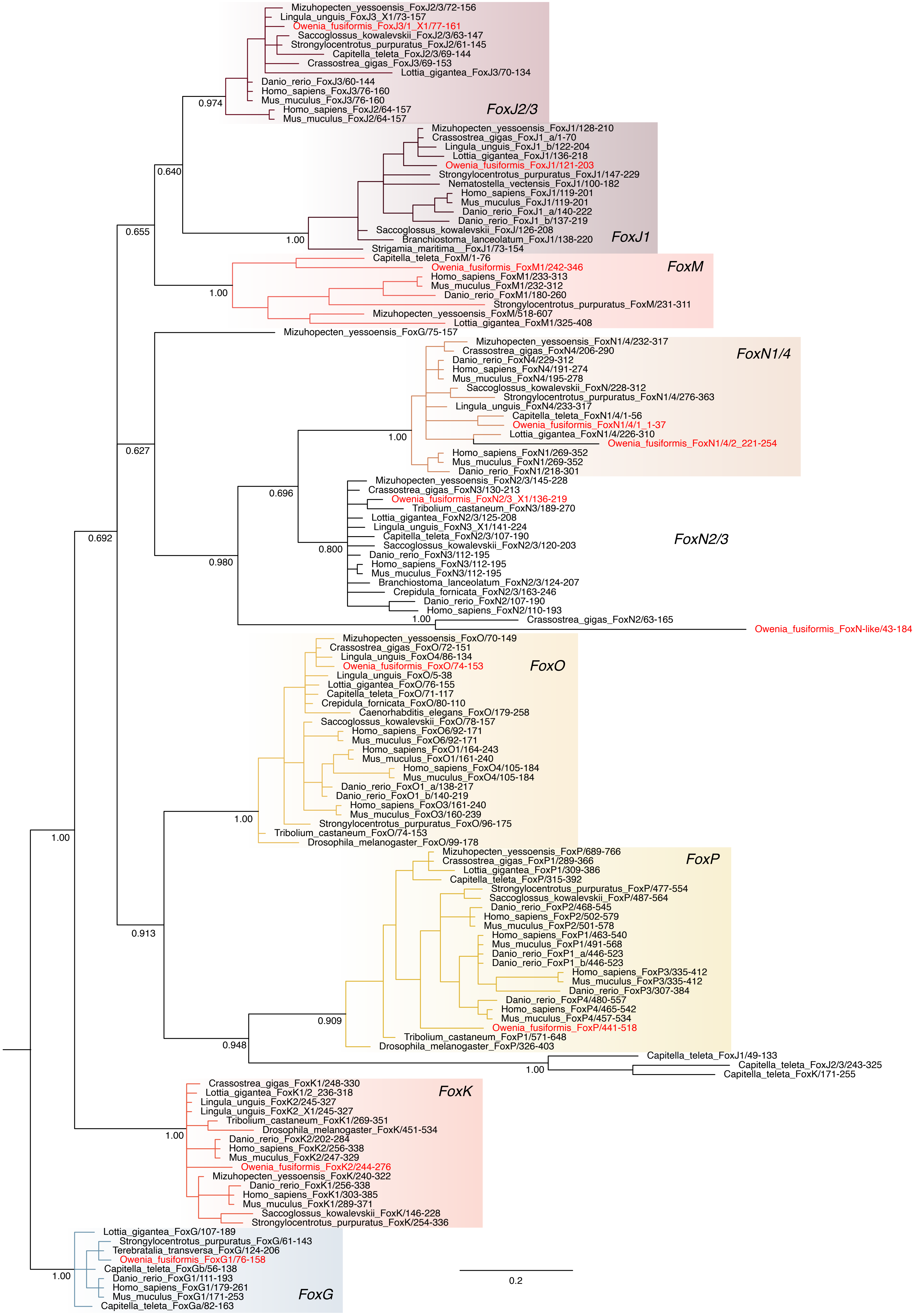

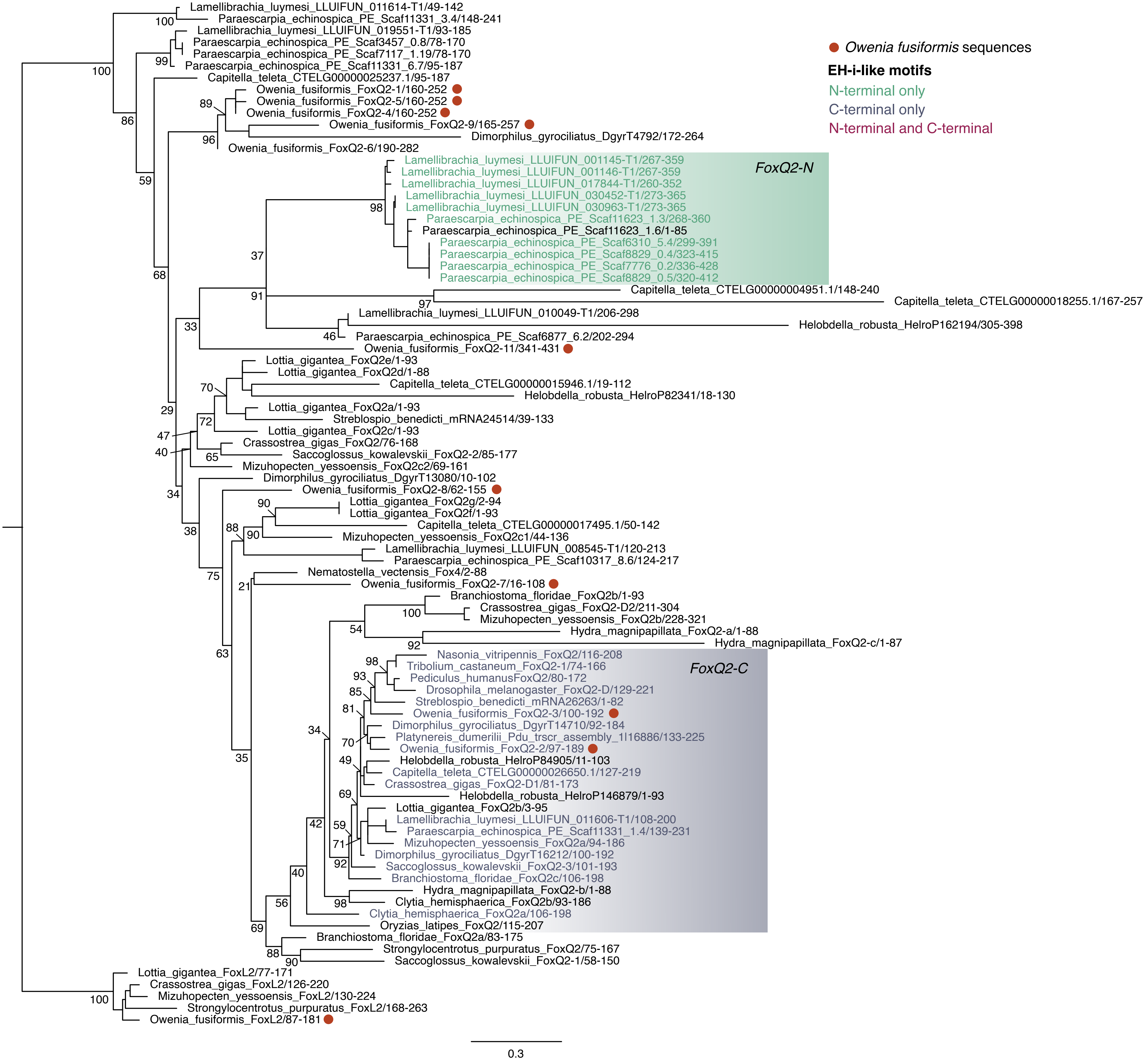

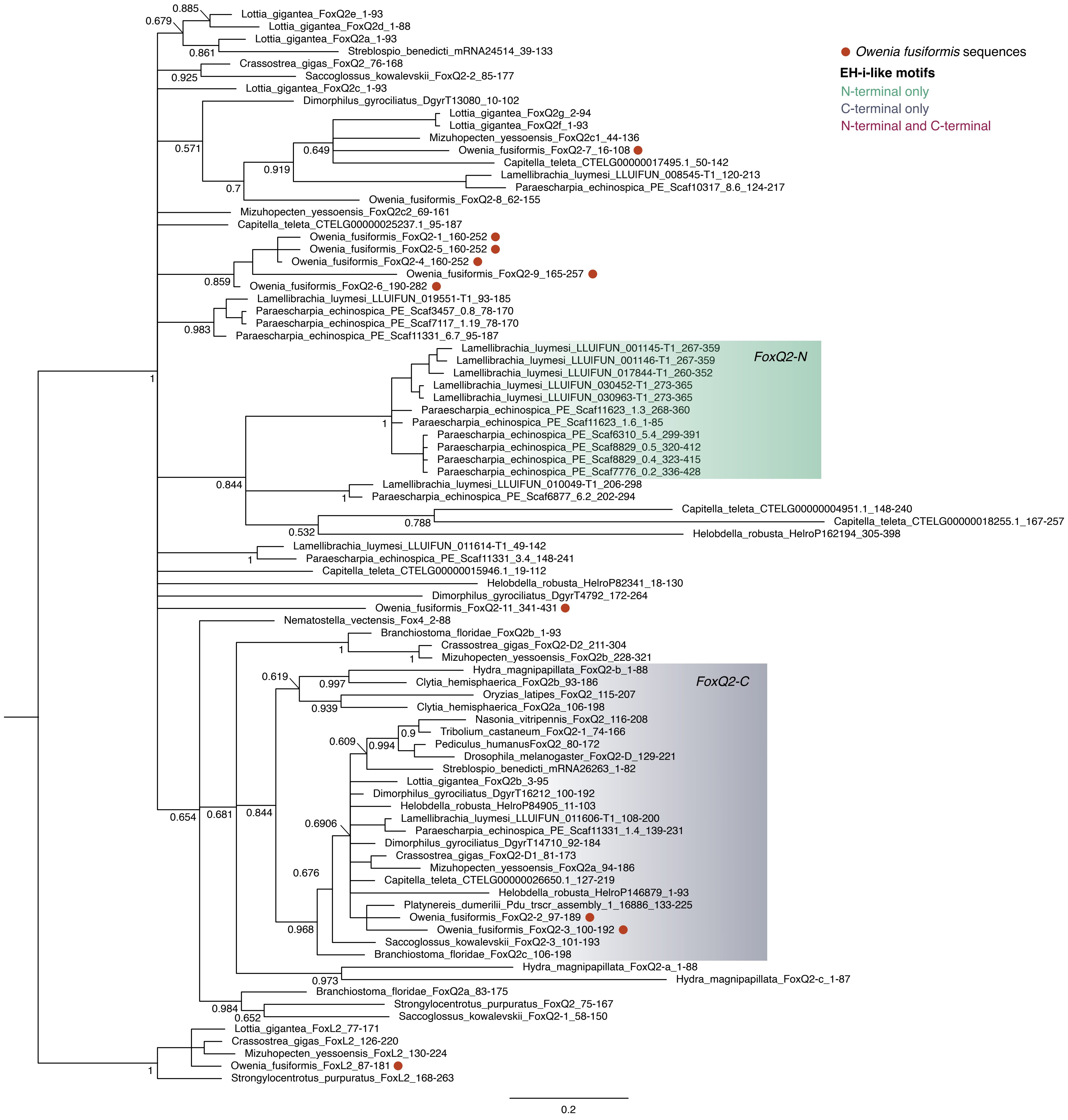

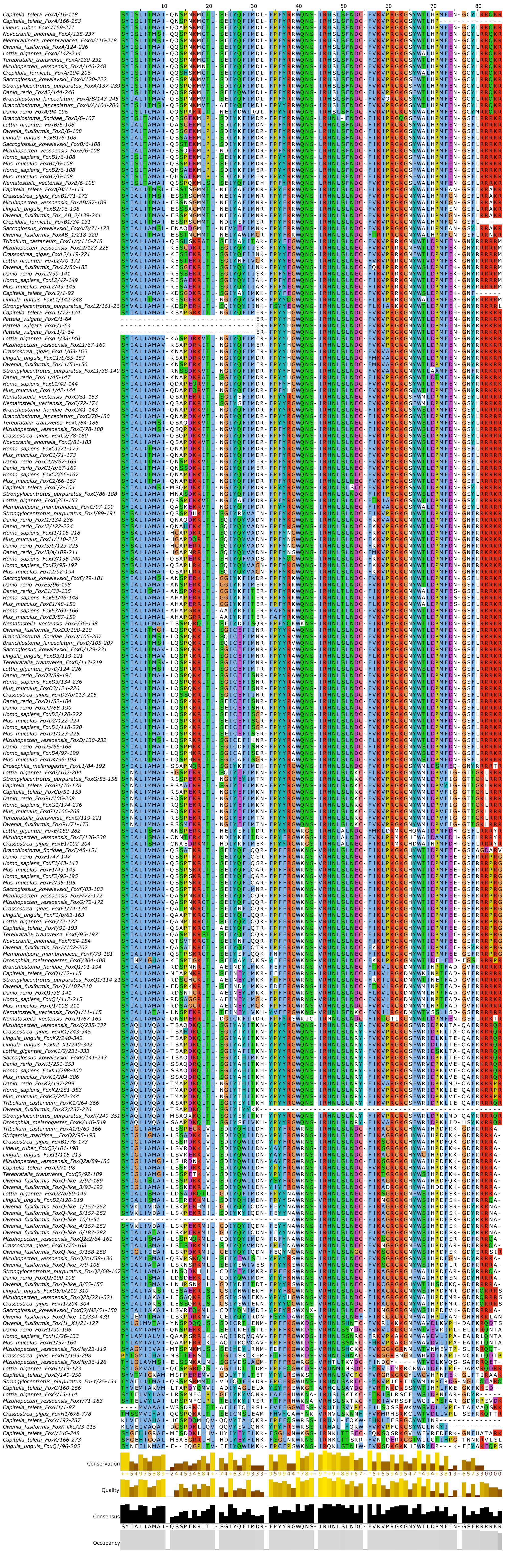

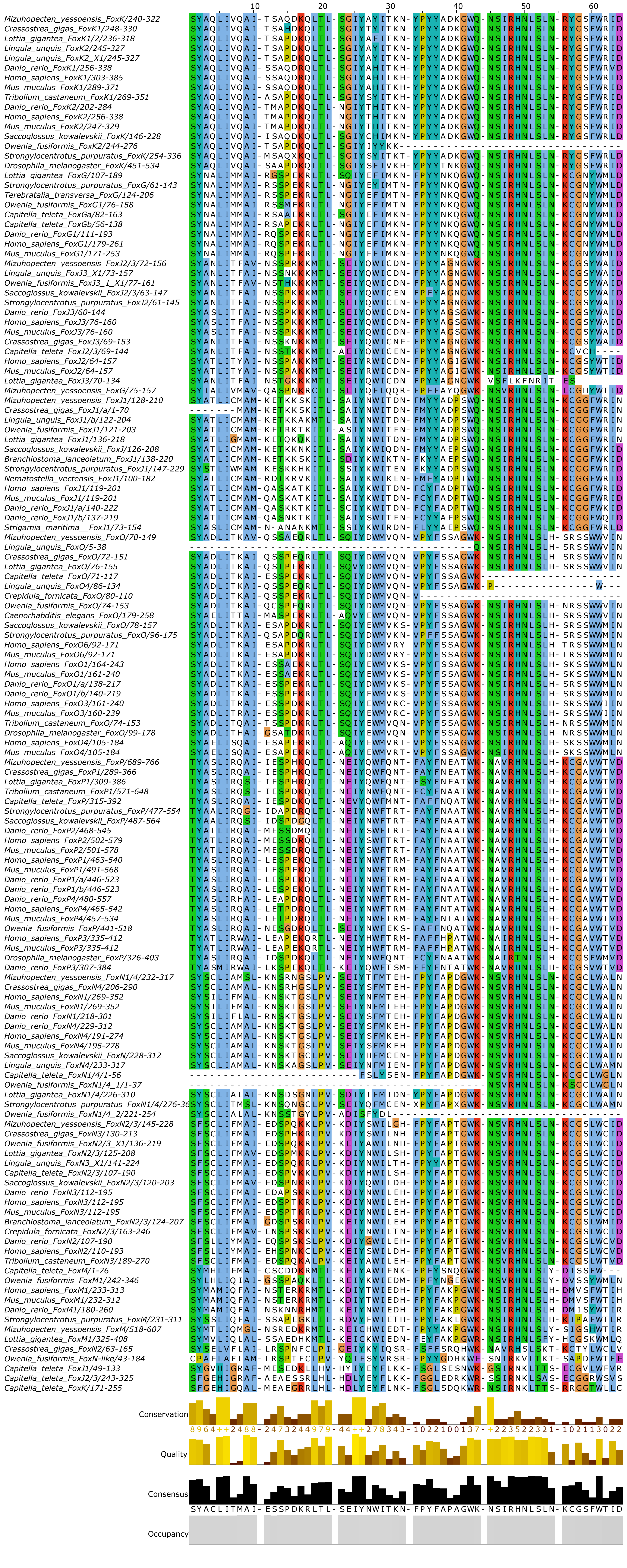

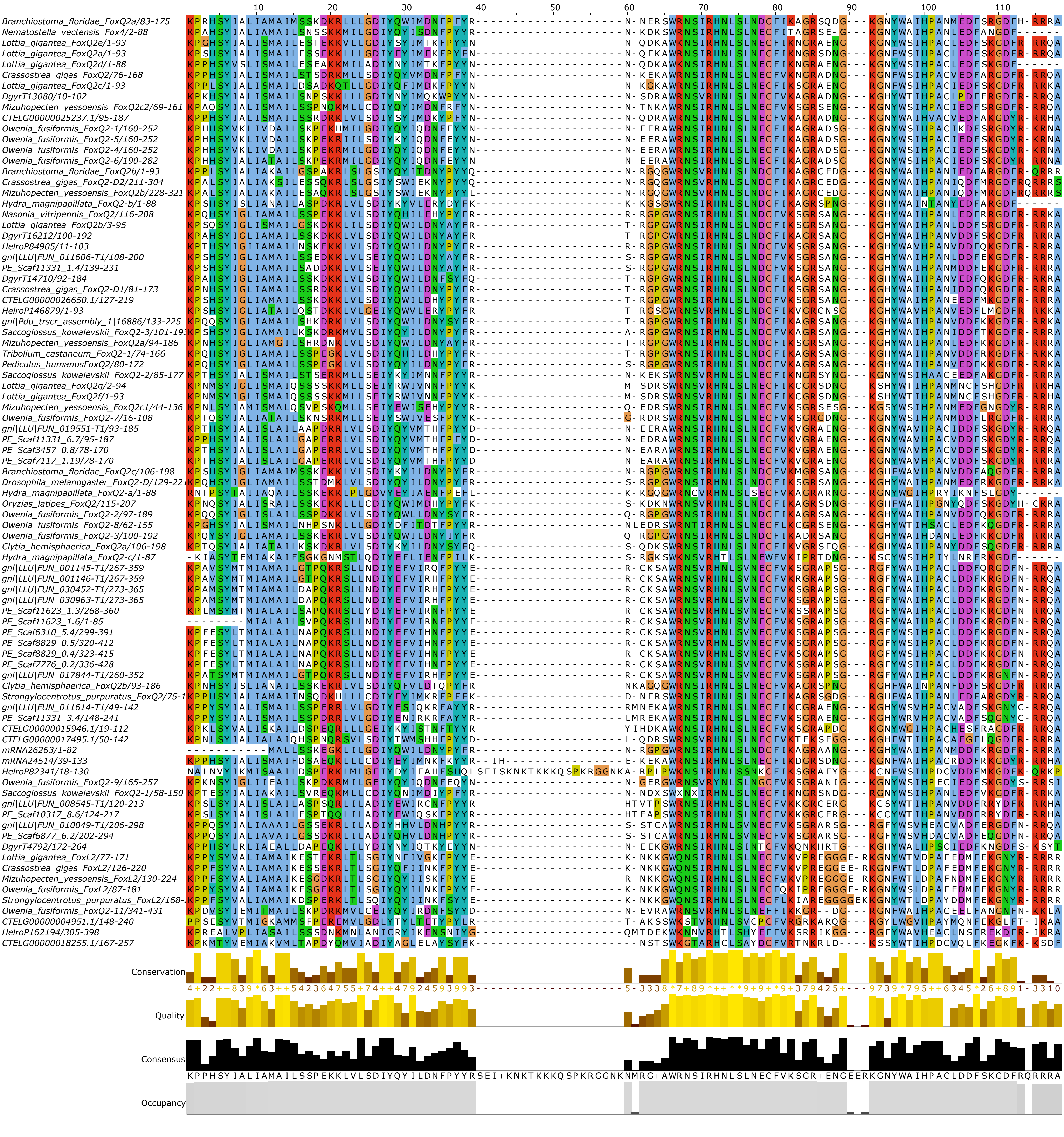
