## Supplementary Tables 1 to 4 for "The *Fox* gene repertoire in the annelid *Owenia fusiformis* reveals multiple expansions of the *foxQ2* class in Spiralia"

**Supplementary Table 1. The expression of *Fox* genes in Spiralia**

| Clade | Genes | Phyla | Species | Gene expression domain | Inferred roles | Reference |
| --- | --- | --- | --- | --- | --- | --- |
| I | FoxA | Annelida | *Hydroides elegans* | *(two paralogs; expression is mostly overlapping)* Blastula: vegetal pole  Gastrula: blastopore, anterior lip of the blastopore | Gut formation and mesoderm development | Arenas-Mena, César. "Embryonic expression of HeFoxA1 and HeFoxA2 in an indirectly developing polychaete." Development genes and evolution 216.11 (2006): 727-736. |
|  |  |  | *Capitella teleta* | Blastula: vegetal blastomeres  Gastrula: blastopore (anterior and posterior).  Larva: foregut, hindgut | Involved in foregut formation. | Boyle, M. J. & Seaver, E. C. Developmental expression of foxA and gata genes during gut formation in the polychaete annelid, Capitella sp. I. Evol. Dev. 10, 89-105 (2008) |
|  |  |  | *Chaetopterus variopedatus* | Blastula: presumptive endoderm  Gastrula: Vegetal cells  Larvae: foregut and hindgut domains | Gut formation | Boyle, Michael J., and Elaine C. Seaver. "Expression of FoxA and GATA transcription factors correlates with regionalized gut development in two lophotrochozoan marine worms: Chaetopterus (Annelida) and Themiste lageniformis (Sipuncula)." EvoDevo 1.1 (2010): 1-18. |
|  |  |  | *Themiste lageniformis* | Blastula: presumptive endoderm Gastrula: Vegetal cells Larvae: foregut and hindgut domains | Gut formation | Boyle, Michael J., and Elaine C. Seaver. "Expression of FoxA and GATA transcription factors correlates with regionalized gut development in two lophotrochozoan marine worms: Chaetopterus (Annelida) and Themiste lageniformis (Sipuncula)." EvoDevo 1.1 (2010): 1-18. |
|  |  |  | *Platynereis dumerilii* | Cleavage: Three large blastomeres in quadrants A, C and B Gastrulation: Early protochophore: Stomodeal plate Middle trochophore: Two bilaterally symmetric patches at the ventral episphere and two cells posterior to the prototroch at the ventrolateral sides of the hyposphere  Metatrochophore: Foregut primordium | Involved in foregut formation Primary role in gastrulation morphogenesis Marker of endoderm formation | Kostyuchenko, Roman P., et al. "FoxA expression pattern in two polychaete species, Alitta virens and Platynereis dumerilii: examination of the conserved key regulator of the gut development from cleavage through larval life, postlarval growth, and regeneration." Developmental Dynamics 248.8 (2019): 728-743. |
|  |  |  | *Alitta virens* | Early trochophore: Individual cells around the blastopore and stomodeal plate Mid-trochophore: Two bilaterally symmetric patches at the ventral episphere and two cells posterior to the prototroch at the ventrolateral sides of the hyposphere  Metatrochophore: Foregut primordium. | Involved in foregut formation Primary role in gastrulation morphogenesis  Marker of endoderm formation | Kostyuchenko, Roman P., et al. "FoxA expression pattern in two polychaete species, Alitta virens and Platynereis dumerilii: examination of the conserved key regulator of the gut development from cleavage through larval life, postlarval growth, and regeneration." Developmental Dynamics 248.8 (2019): 728-743. |
|  |  |  | *Helobdella austinensis* | *(two paralogs)* Cleavage: absents Organogenesis: Expressed in the prostomium and later in the proboscis. | Involved in foregut formation. | Kwak, Hee‐Jin, et al. "Temporal and spatial expression of the Fox gene family in the Leech Helobdella austinensis." Journal of Experimental Zoology Part B: Molecular and Developmental Evolution 330.6-7 (2018): 341-350. |
|  |  |  | *Owenia fusiformis* | Blastula: vegetal macromeres  Gastrula: endoderm, anterior lip blastopore  Larvae: mouth, midgut | Endodermal marker  Foregut marker | Martín-Durán, José M., et al. "The developmental basis for the recurrent evolution of deuterostomy and protostomy." Nature ecology & evolution 1.1 (2016): 1-10.  APA |
|  |  | Mollusca | *Patella vulgata* | Cleavage: faint expression in 3A, 3B and 3C  Blastula: endodermal derivatives, except for 4D.  Gastrula: horseshoe pattern, Ectomesoderm,  Larvae: whole anterior mesodermal field | Gut formation Mesoderm formation  Role in the establishment of anterior inductive patterning | Lartillot N, Le Gouar M, Adoutte A (2002) Expression patterns of fork headand goosecoid homologues in the mollusc Patella vulgata supports the ancestry of the anterior mesendoderm across Bilateria. Dev Genes Evol 212:551–561 |
|  |  |  | *Crepidula fornicata* | Cleavage: All A-D macromeres in regions adjacent to the nuclei  Gastrula: Cells of the blastopore lip Larvae: Developing mouth and esophagus | Endodermal marker  Foregut marker | Perry, K. J., Lyons, D. C., Truchado‐Garcia, M., Fischer, A. H., Helfrich, L. W., Johansson, K. B., ... & Henry, J. Q. (2015). Deployment of regulatory genes during gastrulation and germ layer specification in a model spiralian mollusc Crepidula. Developmental Dynamics, 244(10), 1215-1248.  ISO 690 |
|  |  | Brachiopoda | *Novocrania anomala* | Blastula:  prospective endoderm in the gastral plate  Gastrula: archenteron wall Larvae: gut, mouth and ventral ectoderm | Endodermal marker  Foregut marker | Martín-Durán, José M., et al. "The developmental basis for the recurrent evolution of deuterostomy and protostomy." Nature ecology & evolution 1.1 (2016): 1-10.  APA |
|  |  |  | *Terebratalia transversa* | Blastula: prospective endoderm in the gastral plate  Gastrula: archenteron wall Larvae: gut, mouth and anteroventral ectoderm | Endodermal marker  Foregut marker | Martín-Durán, José M., et al. "The developmental basis for the recurrent evolution of deuterostomy and protostomy." Nature ecology & evolution 1.1 (2016): 1-10.  APA |
|  |  | Platyhelminth | *Schmidtea mediterranea* | *(two paralogs)* Adult: FoxA-1 expressed inn the pharynx and its progenitors and FoxA-2 expressed in a dotted pattern all along the animal body, which could be neurons and/or epidermal cells. | Not determined | Pascual-Carreras, E., Herrera-Úbeda, C., Rosselló, M., Coronel-Córdoba, P., Garcia-Fernàndez, J., Saló, E., & Adell, T. (2021). Analysis of Fox genes in Schmidtea mediterranea reveals new families and a conserved role of Smed-foxO in controlling cell death. Scientific reports, 11(1), 1-18. |
|  |  | Nemertea | *Lineus ruber* | Blastula: absent Gastrula: sbsent Larvae: mouth and pharynx  Juvenile: mouth, inner head, ventral side and posterior end of the endoderm | Endodermal marker  Foregut marker | Martín-Durán, J. M., Vellutini, B. C., & Hejnol, A. (2015). Evolution and development of the adelphophagic, intracapsular Schmidt’s larva of the nemertean Lineus ruber. *Evodevo*, *6*(1), 1-18. |
|  | FoxAB | Annelida | *Capitella teleta* | Cleavage: in a single D-quadrant cell  Early gastrula: outside the blastopore in vegetal micromeres Late gastrula: anterior surface cells encircling the site of stomodeum formation Larvae: in subsurface oral ectoderm sur- rounding the buccal tube | Involved in mouth formation and gut formation  Regulation of ectoderm differentiation | Boyle, Michael J., Emi Yamaguchi, and Elaine C. Seaver. "Molecular conservation of metazoan gut formation: evidence from expression of endomesoderm genes in Capitella teleta (Annelida)." EvoDevo 5.1 (2014): 1-19. |
|  | FoxB | Mollusca | *Crepidula fornicata* | Cleavage: in each of the four quadrants, prominently to regions adjacent to the nuclei of the macromeres Epiboly: in vegetal macromeres and in the progeny of the animal micromeres  Gastrulation: cells located around the blastopore Elongation: mesenchyme and developing head | Ectomesoderm formation  Epithelial-mesenchymal transition at the blastopore lip | S |
|  | FoxC | Phoronida | *Phoronopsis harmeri* | Blastula: posterior mesoderm Late gastrula: posterior mesoderm Larvae: posterior mesoderm | Posterior mesoderm formation | Andrikou, Carmen, and Andreas Hejnol. "FGF signaling acts on different levels of mesoderm development within Spiralia." Development 148.10 (2021): dev196089. |
|  |  | Annelida | *Owenia fusiformis* | Blastula:  absent  Gastrula: absent Elongation: most anterior-lateral mesodermal cells and in a few cells in the anterior dorsal side of the embryo Larvae: posterior pharyngeal mesoderm. | Mesodermal marker | Martín-Durán, José M., et al. "The developmental basis for the recurrent evolution of deuterostomy and protostomy." Nature ecology & evolution 1.1 (2016): 1-10. |
|  |  |  | *Helobdella austinensis* | Cleavage: absent Organogenesis: In prostomium and lateral mesoderm. Later in the ventral muscle and in the proboscis | Foregut formation | Kwak, Hee‐Jin, et al. "Temporal and spatial expression of the Fox gene family in the Leech Helobdella austinensis." Journal of Experimental Zoology Part B: Molecular and Developmental Evolution 330.6-7 (2018): 341-350. |
|  |  |  | *Capitella teleta* | Trochophore:  developing mantle cavities, a bilateral pair of mesodermal cells posterior to the prototroch and a number of ventral mesodermal cells adjacent to the mouth and oesophagus | Posterior mesoderm formation/somatic mesoderm formation Part of the *foxC*-*foxF*-*foxL1*-*foxQ1* cluster for mesoderm formation in metazoa | Shimeld, Sebastian M., et al. "Clustered Fox genes in lophotrochozoans and the evolution of the bilaterian Fox gene cluster." Developmental biology 340.2 (2010): 234-248. |
|  |  | Brachiopod | *Novocrania anomala* | Blastula:  ventral anterior mesoderm Gastrula: ventral anterior mesoderm, , spatially separate from the blastoporal opening  Elongation: ventral anterior mesoderm Larvae: ventral anterior mesoderm | Mesodermal marker | Martín-Durán, José M., et al. "The developmental basis for the recurrent evolution of deuterostomy and protostomy." Nature ecology & evolution 1.1 (2016): 1-10. |
|  |  | Platyhelminth | *Schmidtea mediterranea* | *(two paralogs; expression is overlapping)* Adults: around the pharynx and in the pharynx | Not determined | Pascual-Carreras, E., Herrera-Úbeda, C., Rosselló, M., Coronel-Córdoba, P., Garcia-Fernàndez, J., Saló, E., & Adell, T. (2021). Analysis of Fox genes in Schmidtea mediterranea reveals new families and a conserved role of Smed-foxO in controlling cell death. Scientific reports, 11(1), 1-18. |
|  |  | Mollusca | *Terebratalia transversa* | Gastrula: anterior of the archenteron wall and broadly in the adjacent anterior ectoderm. Elongation:  anterior archenteron wall and anterior ectodermal in the form of  two lateral bands that extend along the animal-vegetal axis. Larvae: ventral anterior and posterior mesoderm. | Mesoderm formation  Evolutionary conserved role for patterning mesoderm at both the anterior and posterior extremities | Passamaneck, Yale J., Andreas Hejnol, and Mark Q. Martindale. "Mesodermal gene expression during the embryonic and larval development of the articulate brachiopod Terebratalia transversa." Evodevo 6.1 (2015): 1-21. |
|  |  |  | *Patella vulgata* | Early trochophore: cells visible in the mantle cavities and cells lying adjacent and anterior to the mouth, under the ectoderm  Late trochophore:: mantle cavities,  mesodermal cells posterior to the prototroch and adjacent to the foot and anterior to the prototroch and adjacent to the oesophagus | Posterior mesoderm formation/somatic mesoderm formation Part of the *foxC*-*foxF*-*foxL1*-*foxQ1* cluster for mesoderm formation in metazoa | Shimeld, Sebastian M., et al. "Clustered Fox genes in lophotrochozoans and the evolution of the bilaterian Fox gene cluster." Developmental biology 340.2 (2010): 234-248. |
|  | FoxD | Annelida | *Platynereis dumerilii* | Adult: ventral somatic muscle markers | Ventral mesoderm marker | Lauri, Antonella, et al. "Development of the annelid axochord: insights into notochord evolution." Science 345.6202 (2014): 1365-1368. |
|  |  | Platyhelminthes | *Schmidtea mediterranea* | Adult: anterior muscle cells | Not determined | Pascual-Carreras, E., Herrera-Úbeda, C., Rosselló, M., Coronel-Córdoba, P., Garcia-Fernàndez, J., Saló, E., & Adell, T. (2021). Analysis of Fox genes in Schmidtea mediterranea reveals new families and a conserved role of Smed-foxO in controlling cell death. Scientific reports, 11(1), 1-18. |
|  |  | Brachiopod | *Terebratalia transversa* | Gastrula: narrow band of cells at the border of the archenteron wall and roof in the radial gastrula and in ectodermal cells at the anterior of the animal Elongation:  narrow band of cells at the border of the archenteron wall and roof in the radial gastrula and broad band of ventral ectodermal expression just anterior of the blastopore Larvae: two bands of the mesoderm in the mantle lobe which converge ventromedially in the pedicle lobe | Ventral mesoderm formation | Passamaneck, Yale J., Andreas Hejnol, and Mark Q. Martindale. "Mesodermal gene expression during the embryonic and larval development of the articulate brachiopod Terebratalia transversa." Evodevo 6.1 (2015): 1-21. |
|  | FoxE |  |  | Not determined |  |  |
|  | FoxF | Phoronida | *Phoronopsis harmeri* | Blastula: absent Late gastrula: anterior mesoderm Larvae: anterior and posterior mesoderm | Anterior and posterior mesoderm formation | Andrikou, Carmen, and Andreas Hejnol. "FGF signaling acts on different levels of mesoderm development within Spiralia." Development 148.10 (2021): dev196089. |
|  |  | Brachiopoda | *Novocrania anomala* | Blastula:  ventral anterior mesoderm Gastrula: ventral anterior mesoderm, , spatially separate from the blastoporal opening  Elongation: ventral anterior mesoderm Larvae: ventral anterior mesoderm | Mesodermal marker | Martín-Durán, José M., et al. "The developmental basis for the recurrent evolution of deuterostomy and protostomy." Nature ecology & evolution 1.1 (2016): 1-10. |
|  |  | Annelida | *Owenia fusiformis* | Blastula and gastrula: most anterior-lateral mesodermal cells and in a few cells in the anterior dorsal side of the embryo Larvae: posterior pharyngeal mesoderm | Mesodermal marker | Martín-Durán, José M., et al. "The developmental basis for the recurrent evolution of deuterostomy and protostomy." Nature ecology & evolution 1.1 (2016): 1-10. |
|  |  |  | *Capitella teleta* | Trochophore:  bilateral expression pattern in the brain,  foregut and posterior ventro-lateral mesoderm | Posterior mesoderm formation/visceral mesoderm formation Part of the *foxC*-*foxF*-*foxL1*-*foxQ1* cluster for mesoderm formation in metazoa | Shimeld, Sebastian M., et al. "Clustered Fox genes in lophotrochozoans and the evolution of the bilaterian Fox gene cluster." Developmental biology 340.2 (2010): 234-248. |
|  |  | Platyhelminthes | *Schmidtea mediterranea* | Adult: cells in the margin of the head and  muscle cells in the lateral dorsal part of animal, between the pharynx and the margin of the organism | Not determined | Pascual-Carreras, E., Herrera-Úbeda, C., Rosselló, M., Coronel-Córdoba, P., Garcia-Fernàndez, J., Saló, E., & Adell, T. (2021). Analysis of Fox genes in Schmidtea mediterranea reveals new families and a conserved role of Smed-foxO in controlling cell death. Scientific reports, 11(1), 1-18. |
|  |  | Brachiopoda | *Terebratalia transversa* | Gastrula: absent Elongation:  anterior of the archenteron wall Larvae: mesoderm laterally and anteriorly flanking the endoderm in larval stages | Mesoderm formation  Larval endoderm formation | Passamaneck, Yale J., Andreas Hejnol, and Mark Q. Martindale. "Mesodermal gene expression during the embryonic and larval development of the articulate brachiopod Terebratalia transversa." Evodevo 6.1 (2015): 1-21. |
|  |  | Mollusca | *Patella vulgata* | Trochophore: in two bilateral pairs of cells, one level with the mouth and one anterior to the prototroch and their descendants in the lateral endoderm. Later, in the developing foot field | Posterior mesoderm formation/visceral mesoderm formation Part of the *foxC*-*foxF*-*foxL1*-*foxQ1* cluster for mesoderm formation in metazoa | Shimeld, Sebastian M., et al. "Clustered Fox genes in lophotrochozoans and the evolution of the bilaterian Fox gene cluster." Developmental biology 340.2 (2010): 234-248. |
|  | FoxG | Platyhelminthes | *Schmidtea mediterranea* | Adult: in a subset of muscle and neuronal cells all  along the DV margin, in the dorsal midline and in some scattered cells in the dorsal and ventral part. | Not determined | Pascual-Carreras, E., Herrera-Úbeda, C., Rosselló, M., Coronel-Córdoba, P., Garcia-Fernàndez, J., Saló, E., & Adell, T. (2021). Analysis of Fox genes in Schmidtea mediterranea reveals new families and a conserved role of Smed-foxO in controlling cell death. Scientific reports, 11(1), 1-18. |
|  |  | Brachiopoda | *Terebratalia transversa* | Gastrula: two dis tinct domains in the animal cap ectoderm of the early radial gastrula Elongation: Two additional, more lateral expression domains  Larvae:  region of the transverse ciliated band and later in a ‘U’- shape domain that borders the anterior ventral ectoderm | Anterior patterning  Ciliary band formation | Santagata, Scott, et al. "Development of the larval anterior neurogenic domains of Terebratalia transversa (Brachiopoda) provides insights into the diversification of larval apical organs and the spiralian nervous system." Evodevo 3.1 (2012): 1-21. |
|  | FoxH | Annelida | *Owenia fusiformis* | Blastula: Uniquely in the D-quadrant organiser cell  Gastrula: Mesoderm precursors | Organizing activity | Seudre, O., Carrillo-Baltodano, A. M., Liang, Y., & Martín-Durán, J. M. (2021). ERK1/2 is an ancestral organising signal in spiral cleavage. *bioRxiv*. |
|  | FoxI |  |  | Not determined | Not determined |  |
|  | FoxJ1 | Annelida | *Platynereis dumerilii* | Trochophore:  Ampullary cells, crescent cells and prototroch  Adult: Ciliated cells | Formation of the apical organ  Formation of the cilia | Marlow, H., Tosches, M. A., Tomer, R., Steinmetz, P. R., Lauri, A., Larsson, T., & Arendt, D. (2014). Larval body patterning and apical organs are conserved in animal evolution. BMC biology, 12(1), 1-17. And Pascual-Carreras, E., Herrera-Úbeda, C., Rosselló, M., Coronel-Córdoba, P., Garcia-Fernàndez, J., Saló, E., & Adell, T. (2021). Analysis of Fox genes in Schmidtea mediterranea reveals new families and a conserved role of Smed-foxO in controlling cell death. Scientific reports, 11(1), 1-18. |
|  | FoxJ3 |  |  | Not determined |  |  |
|  | FoxL1 | Annelida | *Owenia fusiformis* | Blastula and gastrula: anterior portion of the mesodermal bands Larvae: lateral–posterior mesoderm of the pharynx. | Mesodermal marker | Martín-Durán, José M., et al. "The developmental basis for the recurrent evolution of deuterostomy and protostomy." Nature ecology & evolution 1.1 (2016): 1-10. |
|  |  | Mollusca | *Patella vulgata* | Blastula:  4 micromere cells,  in each embryonic quadrant. Trochophore:  bilateral pair of mesoderm cells adjacent to the mouth in Later, two small expression domains are visible on either side of the mouth, lying under the ectoderm and adjacent to the epithelium of the oesophagus | Formation of the mesoderm that lines the external surface of the anterior gut | Shimeld, Sebastian M., et al. "Clustered Fox genes in lophotrochozoans and the evolution of the bilaterian Fox gene cluster." Developmental biology 340.2 (2010): 234-248. |
|  |  |  | *Capitella teleta* | Trochophore:  surface cells on the posterior face of the mouth,  bilateral pair of patches on the dorsal–posterior side of the foregut, small domain bordering the left and right lateral–anterior rims of the pharynx pad and at very low levels in the brain and pharynx regions | Formation of the mesoderm that lines the external surface of the anterior gut | Shimeld, Sebastian M., et al. "Clustered Fox genes in lophotrochozoans and the evolution of the bilaterian Fox gene cluster." Developmental biology 340.2 (2010): 234-248. |
|  | FoxL2 | Annelida | *Helobdella austinensis* | Cleavage: absent Organogenesis: exterior germinal plate and later in mesodermal muscle fiber | Mesoderm development during late embryonic stage | Kwak, Hee‐Jin, et al. "Temporal and spatial expression of the Fox gene family in the Leech Helobdella austinensis." Journal of Experimental Zoology Part B: Molecular and Developmental Evolution 330.6-7 (2018): 341-350. |
|  | FoxQ1 | Annelida | *Capitella teleta* | Trochophore: oesophagus in a highly asymmetric pattern between left and right sides of the animal with higher levels of expression on the left side | Anterior gut formation | Shimeld, Sebastian M., et al. "Clustered Fox genes in lophotrochozoans and the evolution of the bilaterian Fox gene cluster." Developmental biology 340.2 (2010): 234-248. |
|  | FoxQ2 | Annelida | *Platynereis dumerilii* | Trochophore:  Upper two thirds of the episphere  (the apical plate) Adult: in differentiated eye cells, some brain progenitors and in ventral nerve cords | Formation of the apical organ | Marlow, H., Tosches, M. A., Tomer, R., Steinmetz, P. R., Lauri, A., Larsson, T., & Arendt, D. (2014). Larval body patterning and apical organs are conserved in animal evolution. BMC biology, 12(1), 1-17. And Pascual-Carreras, E., Herrera-Úbeda, C., Rosselló, M., Coronel-Córdoba, P., Garcia-Fernàndez, J., Saló, E., & Adell, T. (2021). Analysis of Fox genes in Schmidtea mediterranea reveals new families and a conserved role of Smed-foxO in controlling cell death. Scientific reports, 11(1), 1-18. |
|  |  | Brachiopoda | *Terebratalia transversa* | Gastrula: Asymmetric domain shifted toward the presumptive dorsal end of the anterior ectoderm more Elongation: Subset of anterior dorsal ectodermal domain Larvae: anterior dorsal ectoderm of the apical lobe of the larva as well as a few small dorsal and ventral spots of expression | Anterior patterning  Apical tuft formation Apical organ formation | Santagata, Scott, et al. "Development of the larval anterior neurogenic domains of Terebratalia transversa (Brachiopoda) provides insights into the diversification of larval apical organs and the spiralian nervous system." Evodevo 3.1 (2012): 1-21. |
|  |  | Nemertea | *Lineus ruber* | Blastula: absent Gastrula: sbsent Larvae: most anterior region of the cephalic discs and in the proboscis  Juvenile: anterior head, including the proboscis | Apical organ formation | Martín-Durán, J. M., Vellutini, B. C., & Hejnol, A. (2015). Evolution and development of the adelphophagic, intracapsular Schmidt’s larva of the nemertean Lineus ruber. *Evodevo*, *6*(1), 1-18. |
|  | FoxY |  |  | Not determined |  |  |
| II | FoxK | Platyhelminthes | *Schmidtea mediterranea* | *(three paralogs, expression is overlapping )* Adult: ubiquitously and specifically in the CNS | Not determined | Pascual-Carreras, E., Herrera-Úbeda, C., Rosselló, M., Coronel-Córdoba, P., Garcia-Fernàndez, J., Saló, E., & Adell, T. (2021). Analysis of Fox genes in Schmidtea mediterranea reveals new families and a conserved role of Smed-foxO in controlling cell death. Scientific reports, 11(1), 1-18. |
|  | FoxM1 |  |  | Not determined | Not determined |  |
|  | FoxN1/4 |  |  | Not determined | Not determined |  |
|  | FoxN2/3 | Mollusca | *Crepidula fornicata* | Cleavage: faintly and diffusely in cells in each of the four quadrants Epiboly: In the micromere progeny Gastrulation: around the blastopore lip Elongation: entire lip of the blastopore expresses, including the ectomesodermal cells, in scattered mesenchymal cells leaving the lip of the blastopore, in some progeny of 4d  Larvea: in the developing head | Epithelial-mesenchymal transition at the blastopore lip | Osborne, C. C., Perry, K. J., Shankland, M., & Henry, J. Q. (2018). Ectomesoderm and epithelial–mesenchymal transition‐related genes in spiralian development. *Developmental Dynamics*, *247*(10), 1097-1120. |
|  |  | Platyhelminthes | *Schmidtea mediterranea* | *(two paralogs, expression is overlapping )* Adult: ubiquitously in the SNC | Not determined | Pascual-Carreras, E., Herrera-Úbeda, C., Rosselló, M., Coronel-Córdoba, P., Garcia-Fernàndez, J., Saló, E., & Adell, T. (2021). Analysis of Fox genes in Schmidtea mediterranea reveals new families and a conserved role of Smed-foxO in controlling cell death. Scientific reports, 11(1), 1-18. |
|  | FoxO | Annelida | *Helobdella austinensis* | *(two paralogs - overlapping expression)* Cleavage: Ubiquitous  Organogenesis- FoxO1: Teloblasts and germinal band | Involved in cell division process during cleavage and perform a variety of functions throughout segmentation and organogenesis | Kwak, Hee‐Jin, et al. "Temporal and spatial expression of the Fox gene family in the Leech Helobdella austinensis." Journal of Experimental Zoology Part B: Molecular and Developmental Evolution 330.6-7 (2018): 341-350. |
|  |  | Mollusca | *Crepidula fornicata* | Cleavage: diffusely throughout the micromeres Epiboly: cells around the lip of the blastopore Gastrulation: cells around the lip of the blas- topore, including the ectomesodermal progeny of 3a2 and 3b2   and in the mesentoblast Elongation: cells around the blastopore Larvae: head | General role in morphogenetic processes | Osborne, C. C., Perry, K. J., Shankland, M., & Henry, J. Q. (2018). Ectomesoderm and epithelial–mesenchymal transition‐related genes in spiralian development. *Developmental Dynamics*, *247*(10), 1097-1120. |
|  |  | Platyhelminthes | *Schmidtea mediterranea* | Adult: ubiquitously | Role in controlling cell death | Pascual-Carreras, E., Herrera-Úbeda, C., Rosselló, M., Coronel-Córdoba, P., Garcia-Fernàndez, J., Saló, E., & Adell, T. (2021). Analysis of Fox genes in Schmidtea mediterranea reveals new families and a conserved role of Smed-foxO in controlling cell death. Scientific reports, 11(1), 1-18. |
|  | FoxP | Platyhelminthes | *Schmidtea mediterranea* | Adult: specific parenchymal cell, type, pigment cells | Not determined | Pascual-Carreras, E., Herrera-Úbeda, C., Rosselló, M., Coronel-Córdoba, P., Garcia-Fernàndez, J., Saló, E., & Adell, T. (2021). Analysis of Fox genes in Schmidtea mediterranea reveals new families and a conserved role of Smed-foxO in controlling cell death. Scientific reports, 11(1), 1-18. |

### Supplementary Table 2. Genomic characteristics of the Fox genes in *O. fusiformis*

| transcript id | Fox id | CDS Size (bp) | Number of introns |
| --- | --- | --- | --- |
| OFUSG05364.1 | FoxL1 | 2223 | 0 |
| OFUSG08383.1 | FoxM1 | 2170 | 7 |
| OFUSG05701.1 | FoxP | 2042 | 11 |
| OFUSG01726.1 | FoxO | 1779 | 2 |
| OFUSG27049.1 | FoxC | 1572 | 0 |
| OFUSG21828.1 | FoxN2/3 | 1536 | 6 |
| OFUSG26541.1 | FoxN-like | 1512 | 2 |
| OFUSG13716.1 | FoxD3 | 1488 | 0 |
| OFUSG23528.1 | FoxJ3 | 1466 | 10 |
| OFUSG19661.1 | FoxQ2-11 | 1440 | 0 |
| OFUSG04768.1 | FoxF | 1410 | 1 |
| OFUSG21527.1 | FoxA | 1374 | 0 |
| OFUSG09642.1 | FoxAB-a | 1353 | 0 |
| OFUSG10613.1 | FoxJ1 | 1323 | 1 |
| OFUSG03084.1 | FoxQ2-6 | 1296 | 1 |
| OFUSG09682.1 | FoxQ2-2 | 1269 | 0 |
| OFUSG03081.1 | FoxQ2-1 | 1222 | 0 |
| OFUSG03083.1 | FoxQ2-4 | 1182 | 0 |
| OFUSG03082.1 | FoxQ2-5 | 1182 | 0 |
| OFUSG09693.1 | FoxAB-b | 1134 | 2 |
| OFUSG09683.1 | FoxQ2-3 | 1091 | 0 |
| OFUSG23111.1 | FoxQ1 | 1086 | 0 |
| OFUSG25241.1 | FoxG1 | 1081 | 0 |
| OFUSG20993.1 | FoxQ2-9 | 1017 | 0 |
| OFUSG04867.1 | FoxH1 | 1005 | 0 |
| OFUSG09476.1 | FoxB | 888 | 0 |
| OFUSG23682.1 | FoxL2 | 872 | 2 |
| OFUSG10014.1 | FoxK | 855 | 2 |
| OFUSG08948.1 | FoxQ2-7 | 816 | 0 |
| OFUSG25429.1 | FoxN1/4-a | 792 | 1 |
| OFUSG09684.1 | FoxQ2-8 | 675 | 0 |
| OFUSG24642.1 | FoxQ2-10 | 619 | 0 |
| OFUSG25458.1 | FoxY | 540 | 4 |
| OFUSG25430.1 | FoxN1/4-b | 339 | 0 |

### Supplementary Table 3. Timing of sample collection in four spiralian species

| Species | Developmental stage | Time point | Number of replicates | Source |
| --- | --- | --- | --- | --- |
| *C. teleta* | oocyte | 0 hpf^1^ | 2 | Martin Duran Lab |
|  | zygote | ~ 0.5 hpf | 2 |  |
|  | 2-cell | ~ 1h45 mpf | 2 |  |
|  | 4-cell | ~ 2 hpf | 2 |  |
|  | 8-cell | ~ 4h10 mpf^2^ | 2 |  |
|  | 16-cell | ~ 6h10 mpf | 2 |  |
|  | 32-cell | ~ 8h10 mpf | 2 |  |
|  | blastula | ~ 10h25 mpf | 2 |  |
|  | gastrula | ~ 30h45 mpf | 2 |  |
|  | st4 | ~ 2 dpf^3^ | 2 |  |
|  | st5 | ~ 3 dpf | 2 |  |
|  | St7 | ~ 5 dpf | 2 |  |
| *C. gigas* | oocyte | 0 hpf | 1 | Wang Jun et al. 2012. doi: 10.1038/nature11413. |
|  | 2-cell | 1h20 mpf | 1 |  |
|  | 4-cell | 1h32 mpf | 1 |  |
|  | early morula | 2h25 mpf | 1 |  |
|  | morula | 3.5 hpf | 1 |  |
|  | blastula | 4.5 hpf | 1 |  |
|  | rotary movement | 5.5 hpf | 1 |  |
|  | free swimming | 6.5 hpf | 1 |  |
|  | early gastrula (e. gastrula) | 7.5 hpf | 1 |  |
|  | gastrula | 8.5 hpf | 1 |  |
|  | trocophore | 9.5 hpf | 5 |  |
|  | early D-shape larva | 15.5 hpf | 2 |  |
|  | D-shape larva (D.larva) | 17.5 hpf | 7 |  |
|  | early U-shape larva | 5dpf | 2 |  |
|  | U-shape larva (U.larva) | 10dpf | 6 |  |
|  | Late U-shape larva | 14dpf | 2 |  |
|  | pediveliger | 18dpf | 2 |  |
|  | spat | 22dpf | 1 |  |
|  | juvenile | 215dpf | 1 |  |
| *M. yessoensis* | 2-8cell | 6 hpf | 1 | Wang S et al. 2017. doi: 10.1038/s41559-017-0120. |
|  | Blastula | 18 hpf | 1 |  |
|  | gastrula | 28 hpf | 1 |  |
|  | trocophore | 47 hpf | 1 |  |
|  | D-shape larva | 70 hpf | 1 |  |
|  | pediveliger | 26dpf | 1 |  |
|  | juvenile | 30dpf | 1 |  |
| *O. fusiformis* | oocyte | 0 hpf | 2 | Martin Duran Lab |
|  | zygote | 0.5 hpf | 2 |  |
|  | 2-cell | 1 hpf | 2 |  |
|  | 4-cell | 1.5hf | 2 |  |
|  | 8-cell | 2 hpf | 2 |  |
|  | 16-cell | 3 hpf | 2 |  |
|  | 32-cell | 4 hpf | 2 |  |
|  | blastula | 5 hpf | 2 |  |
|  | gastrula | 9 hpf | 2 |  |
|  | Elongation (elong) | 13 hpf | 2 |  |
|  | Early larva (e. larva) | 17 hpf | 2 |  |
|  | Mitraria Larva | 27 hpf | 2 |  |
|  | Co mpetent larva (c. larva) | 3 wpf | 2 |  |
|  | Juvenile | 1 hpm4 | 1 |  |

^1^ hpf: hour post fertilisation

^2^ mpf: minute post fertilisation

^3^ dpf: day post fertilisation

^4^ hpm: hour post metamorphosis

### Supplementary Table 4. Stage specific RNA-seq data (in TPM) for the *foxQ2* paralogs in *C. teleta*

| Gene | Transcript ID | oocyte_rep1 | oocyte_rep2 | zygote_rep1 | zygote_rep2 | 2cells_rep1 | 2cells_rep2 | 4cells_rep1 |
| --- | --- | --- | --- | --- | --- | --- | --- | --- |
| FoxQ2-1 | CTELG00000004951.1 | 626.45 | 396.52 | 418.56 | 333.22 | 247.13 | 458.14 | 293.41 |
| FoxQ2-2 | CTELG00000015946.1 | 24.48 | 44.50 | 173.64 | 153.10 | 73.82 | 74.11 | 49.52 |
| FoxQ2-3 | CTELG00000017495.1 | 0.00 | 0.00 | 0.00 | 0.00 | 1.26 | 0.00 | 0.00 |
| FoxQ2-4 | CTELG00000018255.1 | 13,689.07 | 14,073.05 | 49,786.80 | 38,187.13 | 48,411.75 | 47,568.38 | 36,141.02 |
| FoxQ2-5 | CTELG00000025237.1 | 634.26 | 1,116.09 | 295.36 | 219.44 | 204.91 | 104.59 | 190.35 |
| FoxQ2-6 | CTELG00000026650.1 | 8.56 | 15.72 | 8.27 | 18.79 | 2.49 | 33.75 | 1.44 |
| Gene | Transcript ID | 4cells_rep2 | 8cells_rep1 | 8cells_rep2 | 16cells_rep1 | 16cells_rep2 | 32cells_rep1 | 32cells_rep2 |
| FoxQ2-1 | CTELG00000004951.1 | 286.3 | 198.5 | 383.5 | 458.0 | 479.4 | 158.6 | 159.6 |
| FoxQ2-2 | CTELG00000015946.1 | 64.2 | 63.9 | 49.6 | 312.5 | 530.3 | 957.8 | 1,511.1 |
| FoxQ2-3 | CTELG00000017495.1 | 0.0 | 0.0 | 0.0 | 1.1 | 6.9 | 3.3 | 4.5 |
| FoxQ2-4 | CTELG00000018255.1 | 37,925.3 | 19,499.8 | 22,487.0 | 8,013.4 | 7,744.2 | 3,201.0 | 2,591.6 |
| FoxQ2-5 | CTELG00000025237.1 | 124.7 | 219.5 | 104.4 | 139.2 | 142.8 | 146.1 | 74.5 |
| FoxQ2-6 | CTELG00000026650.1 | 1.2 | 15.2 | 19.6 | 12.2 | 47.4 | 104.9 | 117.2 |
